# Investigating the developmental onset of regenerative potential in the annelid *Capitella teleta*

**DOI:** 10.1101/2023.05.12.540607

**Authors:** Alicia A. Boyd, Elaine C. Seaver

## Abstract

An animal’s ability to regrow lost tissues or structures can vary greatly during its life cycle. The annelid *Capitella teleta* exhibits posterior, but not anterior, regeneration as juveniles and adults. In contrast, embryos display only limited replacement of specific tissues. To investigate when during development *C. teleta* becomes capable of regeneration, we assessed the extent to which larvae can regenerate. We hypothesized that larvae exhibit intermediate regeneration potential and demonstrate some features of juvenile regeneration, but do not successfully replace all lost structures. Both anterior and posterior regeneration potential of larvae were evaluated following amputation. Wound sites were analyzed for re-epithelialization, cell proliferation by EdU incorporation, stem cell and differentiation marker expression by in situ hybridization, presence of neurites and muscle fibers by immunohistochemistry and phalloidin staining respectively, and regrowth of structures. Wound healing occurred within 6 hours of amputation for both anterior and posterior amputations. Cell proliferation at both wound sites was observed for up to 7 days following amputation. In addition, the stem cell marker *vasa* was expressed at anterior and posterior wound sites. However, growth of new tissue was observed only in posterior amputations. Neurites from the ventral nerve cord were also observed at posterior wound sites. *De novo ash* expression in the ectoderm of anterior wound sites indicated neuronal cell specification, although the absence of *elav* expression indicated an inability to progress to neuronal differentiation. In rare instances, cilia and eyes reformed. Both amputations induced expanded expression of the myogenesis gene *MyoD* in pre-existing tissues. Our results indicate that amputated larvae complete early, but not late, stages of regeneration, indicating a gradual acquisition of regenerative ability in *C. teleta.* Furthermore, amputated larvae can metamorphose into burrowing juveniles, including those missing brain and anterior sensory structures. To our knowledge, this is the first study to assess regenerative potential of annelid larvae.

## 1. Introduction

Regeneration is the replacement of tissues or structures after injury or amputation. The ability to regenerate lost tissues varies throughout the animal kingdom (Alvarado, 2000). For example, animals such as planarians and hydra can completely reform any lost structure with hardly any restriction (i.e., whole-body regeneration) (Reddien, 2020; Reddy et al., 2020). Other animals have a more restrictive capacity to regenerate and reform only specific structures, such as limbs in crustaceans (Alwes et al., 2016) and amphibians (Brockes, 1997; Byrnes, 1904; Singer, 1954). Finally, some animals, like the leech, display almost no regenerative capabilities following bisection (Özpolat & Bely, 2016). Here, we briefly review differences in regeneration potential across animal life cycles and describe the need to more carefully examine regeneration in pre-adult stages. We highlight annelids as a clade with exceptional regeneration diversity and introduce *Capitella teleta* as a prime candidate to examine how regeneration changes during the life cycle of an indirect developer.

Most published surveys of regeneration ability focus on adult stages, while few studies examine regeneration potential at earlier stages of the life cycle. Several questions emerge when one considers regeneration potential across the life cycle. In animals with a biphasic life cycle, does regeneration ability differ among embryos, larvae, and adults? If so, when in the life cycle is the regeneration program activated – during larval stages or after metamorphosis? Are there conserved trends of increasing or decreasing regeneration abilities across the life cycle? From a developmental perspective, one might expect a trend of decreasing regeneration abilities through the life cycle as embryonic cells become committed to distinct cell types, undergo terminal differentiation, and lose their pluripotency. As a result, there would be progressive loss in the ability to replace tissues (e.g., appendages, nervous system, and heart) (Yun, 2015; Porello et al., 2011; Walters & Zuo, 2013). However, in animals with indirect life cycles, larvae and adults often have distinct tissues, and adult tissue formation requires the retention of progenitors or multipotent stem cells through the larval phase (Peterson et al., 1997). The presence of stem cells in larvae suggests the possibility that animals with indirect development may have greater regeneration potential relative to direct developing species.

The published studies that do examine regeneration in larvae are disproportionately sampled from deuterostomes. From these examples, a complex picture emerges; a single relationship between larval and adult regeneration ability does not explain published observations across species. For example, the *Xenopus* frog (Phipps et al., 2020) has a pattern of regeneration loss similar to what is observed in direct developers: an ability to replace specific tissues (i.e., appendages, heart, brain, lens) as larvae that is lost with maturation (Harrison, 1898). The crinoid, *Antedon rosacea*, is very different such that the larvae exhibit limited ability to regenerate following bisection, but adults are capable of whole-body regeneration (Carnevali, 2006; Carnevali et al., 1993). Furthermore, sea stars exhibit whole-body regeneration during larval, juvenile, and adult stages (Carnevali, 2006). Other animals have a more dynamic change in regenerative ability through their life cycle. *Ciona intestinalis* lacks regenerative abilities as a larva yet exhibits structural regeneration as adults only to progressively lose this ability with advanced age (Jeffery, 2015). A broad understanding of how regeneration ability relates to the life cycle is largely unexplored, and it is critical to sample pre-adult stages in animal taxa with demonstrated regeneration abilities as adults, particularly from protostome lineages.

Annelids are segmented worms that have a wide range of regenerative abilities as adults. Some lack the ability to regenerate following transverse amputation, others regenerate either anterior or posterior segments only, or even undergo whole-body regeneration (Bely, 2006). A systematic review of annelid regenerative ability revealed that anterior and posterior regeneration are widespread across the phylum, alongside numerous examples of loss of regeneration (Bely, 2006). Furthermore, it is likely that both anterior and posterior regeneration are ancestral for adult annelids (Zattara & Bely, 2016). Although some studies have characterized regeneration during juvenile stages (del Olmo et al., 2021; Kostyuchenko, 2022; Planques et al., 2019), to our knowledge, a broad assessment of larval regenerative potential is lacking for annelids.

*Capitella teleta* BLAKE ET AL 2009 is an annelid capable of posterior regeneration and has an indirect life cycle (Seaver, 2022). Previous studies hint at changes in the ability of *C. teleta* to replace lost structures across its life cycle. Juvenile and adult worms regenerate following bisection along the main body axis (de Jong & Seaver, 2016; Giani et al., 2011). In contrast, early-stage embryos do not regulate following deletion of individual precursor cells (Pernet et al., 2012) (apart from eye and germline precursor cells (Dannenberg & Seaver, 2018; Yamaguchi et al., 2016)). It is unknown when *C. teleta* gains its regenerative ability, or to what degree larvae might be capable of regeneration.

Following *C. teleta* embryogenesis, a non-feeding lecithotrophic larva forms. Lecithotrophic larvae often have ‘abbreviated developmental times’ (Vickery et al., 2001), which characterizes the *C. teleta* larval phase of approximately 5 – 6 days in duration. Larvae are characterized by trochal ciliated bands that divide the body into three regions along the anterior–posterior axis: the head, the segmented trunk, and the pygidium. The prototroch separates the head from the trunk and the telotroch separates the trunk from the pygidium (Fig 1A, B) (Seaver et al., 2005). These two ciliary bands are composed of long cilia. A third ciliary band, the neurotroch, runs along the ventral midline of the trunk and is composed of short cilia. Short cilia are also present in the pygidium. Larvae utilize the prototroch and telotroch to swim in the water column, aiding in dispersal.

**Figure 1.**
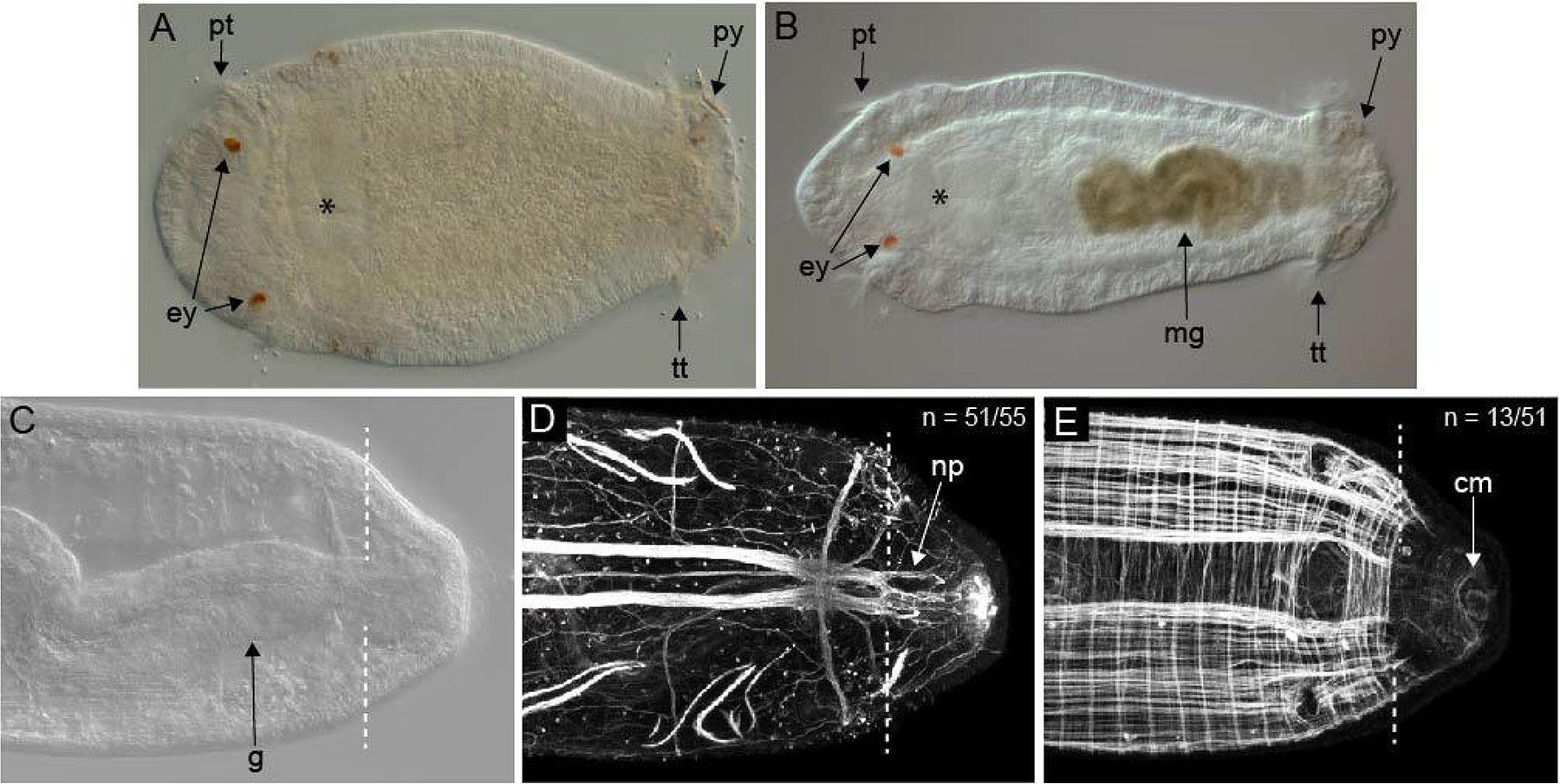
Larvae and regenerating juveniles of *C. teleta*. All images are oriented in ventral view with anterior to the left. **A** Stage 6 larva. **B** Stage 9 larva. **C**, **D**, and **E** are images of the posterior end of a regenerating juvenile 3 days post-amputation (dpa) and are from the same specimen. **C** Differential interference contrast (DIC) image showing regeneration of new tissue in juveniles. **D** Nervous system visualized with anti-acetylated tubulin antibody labeling. **E** Phalloidin staining shows muscle fibers with few fibers in the regenerating tissue. The white dotted lines indicate amputation sites. The asterisk marks the position of the mouth. N denotes the number of animals scored that resemble the image shown. Cm, circular muscle; ey, eyes; g, gut; mg, midgut; np, neural projection; pt, prototroch; py, pygidium; tt, telotroch.

During larval stages, the adult centralized nervous system, musculature, digestive system, and initial systems mature (Meyer et al., 2015). A previously described staging system uses morphological features to identify nine distinct stages of embryonic (Stages 1-3) and larval development (Stages 4-9) (Seaver et al., 2005). *C. teleta* has a centralized nervous system whose development is initiated in late embryogenesis. A ladder-like ventral nerve cord (VNC) positioned along the midline of the trunk forms and is connected to the brain by a pair of circumoral nerves (Meyer et al., 2015; Seaver et al., 2005). The body wall musculature is organized segmentally with circular and longitudinal muscles, and the first muscle fibers appear at the beginning of larval life (Stage 4). Two pigmented eyes appear at Stage 5 and are located lateral and anterior to the prototroch (Meyer et al., 2015; Yamaguchi et al., 2016). Gut organogenesis becomes visibly pronounced between Stages 6–9. The foregut consists of the mouth, buccal cavity, pharynx (morphogenesis in Stage 7), and esophagus (Boyle & Seaver, 2008; see Fig 1A and Fig 1B). In Stages 8–9, the midgut differentiates, becoming olive green and stereotypically coiled (Fig 1B). Stage 6 larvae have approximately 10 segments, generating one segment a day until there are approximately 13 segments present in Stage 9 larvae (Seaver et al., 2005). Larval development concludes when Stage 9 larvae metamorphose into burrowing juvenile worms (Cohen & Pechenik, 1999; Seaver et al., 2005).

We sought to determine when regeneration potential is initiated during the life history of *C. teleta*. We hypothesized that larvae may have limited regeneration potential distinct from juveniles and adults. We anticipated that larvae would display early stages of regeneration without successfully replacing lost tissues and structures. This would suggest that *C. teleta* acquires its regenerative ability gradually through its life cycle. To test our hypothesis, we developed an amputation protocol for larvae to investigate both anterior and posterior regeneration. We characterized larval responses to amputation by making direct comparisons with regeneration events in juveniles. At the wound site, we documented epithelial wound healing, evaluated cell proliferation by EdU incorporation, analyzed changes in neuronal organization with antibody labeling, and visualized musculature with phalloidin staining. We also conducted *in situ* hybridization experiments to analyze expression of the neuronal specification and differentiation gene markers, *ash1* and *elav*, respectively, as well as examine muscle fate and differentiation through analysis of the *MyoD* gene.

## 2. Materials and Methods

### 2.1 Animal Care

As described in previous work (Grassle & Grassle, 1976), a colony of Capitella teleta was maintained in the laboratory at 19°C in glass finger bowls containing filtered sea water (FSW) and a thin layer of mud. Using forceps, larvae were dissected from the brood tubes of healthy females. Larvae from a single brood tube develop synchronously and the number of individuals/brood averaged 100 – 250 individuals. Brood tubes were maintained in FSW until the desired age for amputation or treatment.

### 2.2 Amputations

Stage 6 larvae were used for amputation experiments. Larvae were developmentally staged following a standard staging system (Seaver et al., 2005). Stage 6 is characterized by presence of yolk in the anterior portion of the larva, swimming behavior with a positive phototactic response, and lack of chaetae. Prior to amputation, larvae were incubated in 1:1 MgCl_2_: FSW for 10 minutes to inhibit muscle contraction. Individual larvae were transferred a 60 mm petri dish lid filled with a methyl cellulose solution (0.3 g methyl cellulose in 15 ml filtered sea water) for amputation. Additional methyl cellulose was added as necessary to achieve immobilization. Glass capillary tubes (50 µl, Becton Dickinson and Company) were heated over a flame and pulled into thin, flexible, sharp tips to use as amputation tools. Larvae were amputated with a single continuous, downward motion. Cut sites for anterior amputations and eye amputations were located just posterior to the prototroch, removing the anterior third of the animal (i.e., brain, eyes, and prototroch). Cut sites for posterior amputations were located ∼2/3 of the body length posterior to where the width of the trunk decreases and slopes to the pygidium, effectively removing the posterior growth zone, telotroch and pygidium. Following amputation, each individual was visually inspected for correct cut location and complete separation of the two tissue fragments. Amputated larvae were transferred to a 60 mm petri dish containing 1:1 0.37M MgCl_2_: FSW and incubated for a minimum of 1 hour. Larvae were then washed into FSW containing 60 µg/ml penicillin and 50 µg/ml streptomycin (Sigma-Aldrich Co.) (P/S) and raised at 19°C. P/S FSW was replaced daily, and vitality and swimming behavior were monitored daily. For each amputation experiment, age matched larvae from the same brood tube as experimental animals were raised as uncut controls. In a typical experiment, 40 – 60 individuals from a single brood were amputated. For posterior amputations, amputations were performed on sets of individuals from at least 10 broods, and for anterior amputations, sets of individuals from at least 5 broods were amputated. Both anterior and posterior amputated larvae had high rates of survivorship, ranging from 75% (n= 37/50) to 85% (n=28/33).

Two-week-old juveniles were prepared for amputation by incubation in 1:1 MgCl_2_: FSW for 10 minutes. Individual worms were transferred to a platform of black dissecting wax (American Educational Products, Fort Collins, CO, USA) in the lid of a 35 mm plastic dish for amputation in a drop of 1:1 MgCl_2_: FSW. For eye amputations, worms were amputated immediately anterior to the mouth, removing the eyes and brain. Amputations for posterior regeneration were at the boundary between segment 10 and 11. Individual worms were visually inspected for the correct cut site and successful separation before being transferred to a 60 mm petri dish and incubated in 1:1 MgCl_2_: FSW for a minimum of 1 hour. The juveniles were then washed into FSW and moved to a 60 mm petri dish containing mud and FSW and raised at 19°C for 1 week. For analysis, juvenile worms were sifted from the mud and placed into a petri dish of FSW. Worms were fixed with 4% paraformaldehyde (PFA) in FSW for 30 min before being washed into phosphate-buffered saline (PBS). Fixed juveniles were scored for pigmented eye cells before being processed for antibody labeling.

### 2.3 Cell Proliferation Assay

Larvae were incubated in EdU (Click-iT EdU Alexa Fluor 488 imaging kit – Invitrogen) diluted in FSW to a final concentration of 3 uM at room temperature for 30 minutes. Larvae were then washed into a 1:1 MgCl_2_: FSW solution for 10 minutes and then fixed in 4% PFA for 30 minutes. Afterward, they were rinsed into PBS and incubated for 20 minutes at room temperature in PBS + 0.05% Triton-X-100 with rocking. The specimens were incubated in the commercially provided EdU reaction buffer for 30 minutes, rocking at room temperature. Finally, larvae were washed into PBS and stored at 4°C until further analysis.

### 2.4 EdU Quantification

Quantification of EdU and DAPI stained cells in uncut and amputated larvae was conducted by importing Z-stacks generated from confocal microscopy (see Microscopy) into ImageJ software. Using DAPI and DIC channels to establish boundaries, each image was cropped around the perimeter of the animal. The channels were split and treated separately to optimize cell counting. The threshold for each channel was adjusted manually to minimize background and enhance distinction of individual cells. Afterward, a mask and watershed application were applied to each channel. The resulting image was divided into three equal sections along the anterior-posterior axis using the Montage to Stack application. The modified images were compiled into 20 slice projections (20 µm total) before being counted automatically by Analyze Particles in the Analyze menu. Each 20 µm Z-stack was counted, with the results automatically summarized via section. Manual inspection eliminated any cells that were counted twice by the software. Results were manually compiled by channel and section for each animal, as well as for each time point. A ratio of EdU stained cells to DAPI stained cells was calculated for each time point from the ImageJ cell count summaries. Averages of each body section (anterior, middle, and posterior) for each time point (uncut controls, 6 hpa, 3dpa, and 5dpa) were calculated separately. Ten randomly selected larvae were scored for each time point and a mean value was calculated for each body section across individuals. Each body section of uncut controls was compared with corresponding body sections of amputated animals. One-way weighted ANOVA statistical tests were performed using VassarStats (Lowry, 2023) to test for significance between the control and amputation group of interest.

### 2.5 Antibody labeling and phalloidin staining

Amputated larvae or juveniles were incubated in a 1:1 MgCl_2_: FSW solution for 10 minutes and fixed in 4% paraformaldehyde in FSW for 30 minutes. Animals were washed several times into PBS to remove fixative and then washed into a PBS + Triton solution (PBT) (0.1% Triton-X-100 for larvae and 0.2% Triton-X-100 for juveniles). Animals were transferred to a 3 well glass depression plate and incubated in a solution of 10% heat inactivated normal goat serum in PBT at room temperature with rocking for 45– 60 min. For antibody labeling, the anti-acetylated tubulin antibody (goat anti-mouse, Sigma-Aldrich Co.) was diluted 1:400 in block solution (10% heated treated goat serum in PBT), and incubated with larvae or juveniles at room temperature for 2 hours or overnight at 4°C. Following multiple washes in PBT (4 times in PBT for 30 minutes each), a secondary antibody (goat anti-mouse red 594, Invitrogen) was diluted 1:400 in block solution and added to animals prior to incubation for 2–4 hours rocking at room temperature or overnight at 4°C. The 22C10 antibody (Goat anti-mouse, Developmental Studies Hybridoma bank; 22C10 was deposited to the DSHB by Benzer, S. / Colley, N. (DSHB Hybridoma Product 22C10)) was diluted 1:200 in block solution, added to animals and incubated overnight at 4°C. Following multiple washes in PBT over two hours, the secondary antibody (goat anti-mouse green 488 – Invitrogen) was diluted 1:400 in block solution, added to animals and incubated overnight at 4°C. Following incubation in secondary antibody, larvae were rinsed 2–3 times in PBT and then washed 4 times in PBT for 30 minutes each wash. For phalloidin staining, specimens were incubated in a 1:400 dilution of phalloidin (Alexa Fluor 488 phalloidin, Invitrogen) in PBT. Specimens were incubated on a rocker at room temperature for 45–60 min before being washed twice in PBT for 30 minutes each wash. Animals were cleared in 80% glycerol in PBS with 1 µg/ml Hoechst-33342 (Molecular Probes) overnight prior to imaging.

### 2.6 Whole mount in situ hybridization

Larvae collected at different time points following amputation were incubated in a 1:1 MgCl_2_: FSW solution for 10 minutes and then fixed in 4% paraformaldehyde overnight at 4°C. Fixative was removed by multiple rinses in PBS and then larvae were gradually washed into 100% methanol and stored at -20°C. Whole mount in situ hybridization was performed following a previously published protocol (de Jong & Seaver, 2017). DIG-labeled RNA probes were generated with the MegaScript SP6 or T7 Transcription kit (Invitrogen) according to manufacturer instructions. The following probes and their working concentrations were used: ash1 with 3ng/ul, MyoD with 0.5ng/ul, vasa with 1ng/ul, and elav with 2ng/ul. Animals were hybridized for ∼24 – 48 hours with riboprobe at 65°C. Each RNA probe was visualized by exposure to an anti-digoxigenin-alkaline phosphatase conjugate antibody (Roche) prior to exposure to a color reaction solution containing NBT/BCIP (nitro blue tetrazolium chloride/5-bromo-4-chloro-3-indolyphosphate) and visually monitored for optimal development time. For each sample, uncut control larvae were developed for the same duration as experimental, amputated larvae. Color reactions were terminated following washes of PBT followed by a final fixation in 4% PFA in FSW for 30 min. Tissue was cleared in 80% glycerol in PBS plus 0.1 ug/ml Hoescht to stain nuclei, and then individuals were placed on microscope slides for analysis and imaging. At least two independent repetitions were performed for each gene.

### 2.7 Tissue and Cilia Measurements

To measure tissue formation in amputated larvae, confocal images were analyzed using ImageJ software. Multiple measurements were made on each specimen. The nervous system was measured from the cut site to the position most distal from the cut site that contained neurites. Muscle measurements were taken to the position most distal from the cut site that contained phalloidin-positive fibers. Changes in body width were calculated by making two measurements per specimen; the first measurement was taken at the axial level of the mouth, and the second was just posterior to the cut site. Gut measurements were made from the posterior end of the pigmented midgut to the posterior edge of the posterior ectoderm. For anterior amputations, the muscle and nervous system were measured on 6 hpa (n = 8) and 3 dpa (n = 10) larvae. For posterior amputations, the muscle, nervous system, and changes in body width were measured for 6 hpa (n = 7) and 5 dpa (n = 10). Additionally, the posterior edge of the midgut was used as a landmark for measurements in 5 dpa and Stage 9 larvae at 6 hpa (n = 10). One-way weighted ANOVA statistical tests were performed using the Good Calculators software (Good calculators, 2023) to test for significance between the uncut control and experimental amputation group of interest.

The cilia of larvae were measured using ImageJ software on confocal image projections. Individual labeled cilia of the prototroch, neurotroch, telotroch, and pygidium were traced. Lines resulting from the trace were then measured using the ‘measure’ tool found under ‘Analyze.’ At least three cilia for each ciliary band type were measured in a representative individual, and three individuals were measured, resulting in a total of at least nine measurements for each of the prototroch, telotroch, neurotroch and pygidium ciliary lengths. The range of cilia lengths across the nine measurements is reported (see Results).

### 2.8 Metamorphosis assays

Metamorphosis is rapidly induced in C. teleta by exposure to vitamin B supplements (Burns et al., 2014). A single multivitamin (Centrum Men Multivitamin) was dissolved in 200 ml FSW and stored at room temperature. To test whether amputated larvae are capable of successful metamorphosis, uncut, anteriorly amputated, and posteriorly amputated larvae were exposed to the multivitamin solution containing vitamin B. Larvae were placed in a 30mm petri dish in FSW. Most of the FSW was removed from the dish and replaced with 3 ml of the multi-vitamin solution. Animals were incubated in the vitamin solution for 20 – 30 minutes. The multi-vitamin solution was removed by a minimum of three washes with FSW (2 ml/wash), and then animals were individually scored for settlement and metamorphosis. Successful settlement and metamorphosis were defined as cessation of swimming behavior and initiation of burrowing, loss of the prototroch and telotroch ciliary bands, and elongation of the body.

### 2.9 Microscopy and Imaging Analysis

Larvae were imaged with a Zeiss 710 confocal microscope using 4x bidirectional and unidirectional scan settings for the green and red lasers. Z-stacks were compiled using ImageJ software (Schindelin et al., 2012). Differential interference contrast microscopy images were captured on a Zeiss AxioSkop II motplus compound microscope (Zeiss) coupled with a SPOT FLEX digital camera (Diagnostic Instruments Inc). Multiple focal planes were merged for DIC images using Helicon Focus software (Helicon Focus 7). Images were cropped and processed for brightness and contrast with Photoshop (Adobe Photoshop 2020) or Illustrator (Adobe Illustrator 2020). Figures were composed in Adobe Illustrator (Adobe Illustrator 2020).

## 3. Results

### 3.1 Characteristics of successful regeneration in C. teleta

To characterize larval regenerative potential, we looked to the juvenile posterior regeneration program in *C. teleta* to define features of successful regeneration. *C. teleta* juveniles maintain three conserved sequential stages of posterior regeneration: wound healing, cell proliferation, and differentiation of new tissues (de Jong & Seaver, 2016). Following amputation, juveniles wound heal in 4–6 hours and thereafter form a blastema (Fig 1C) (de Jong & Seaver, 2016). Neuronal projections extend from the VNC into the wound site as early as 2 days post amputation (dpa) (Fig 1D; see also de Jong & Seaver, 2016). In regenerating juveniles, the thickness of the connective nerve decreases dramatically at the point of amputation, with multiple thin neural projections extending from the old VNC tissue into the wound tissue and thereafter into the blastema/new tissue. This change in VNC thickness indicates the position of the cut site. Localized cell proliferation is also observed at the wound site by 2 dpa and persists for several days as new segments form (de Jong & Seaver, 2016). We observed the density of circular and longitudinal muscle fibers decrease abruptly at the wound site at 3 dpa (Fig 1E). Circular and longitudinal muscle fibers are absent between the posterior boundary of the old tissue and phalloidin-positive fibers at the posterior end of the animal. The decrease in muscle fibers spatially align with the decrease in thickness of the connective nerves of the VNC. After approximately 7 days, additional segments are formed that contain organized ganglia, peripheral nerves, circular and longitudinal muscles, digestive tissue and other differentiated tissues (de Jong & Seaver, 2016).

### 3.2 Amputation responses in larvae and subsequent development

To assess the regenerative potential of *C. teleta* larvae, sites were selected for both anterior and posterior amputations that removed approximately 1/3 of the body (Fig 2A). Initial analysis focused on wound healing, which is identified by presence of a continuous epithelial layer covering the wound site and results from contraction of the severed edges of the body wall. To assess wound healing in amputated larvae, we analyzed Stage 6 larvae fixed at 2 hours post amputation (hpa), 3 hpa, 5 hpa, and 6 hpa for presence of a continuous epithelial layer covering the wound site. Following anterior amputations, we observed consistent wound healing 6 hpa (Fig 2B) (n = 54/54). For posterior amputations, we observed wound healing as early as 2 hpa (n = 3/15); however, in most cases successful wound healing occurred around 6 hpa (Fig 2C) (n = 48/51). In posterior amputations performed on Stage 9 larvae, successful wound healing also occurred by 6 hpa (Fig 2D). Presence of tissue between the posterior edge of the pigmented midgut and the posterior epidermis of most animals indicated wound healing of the mesodermal layer by 6 hpa.

**Figure 2.**
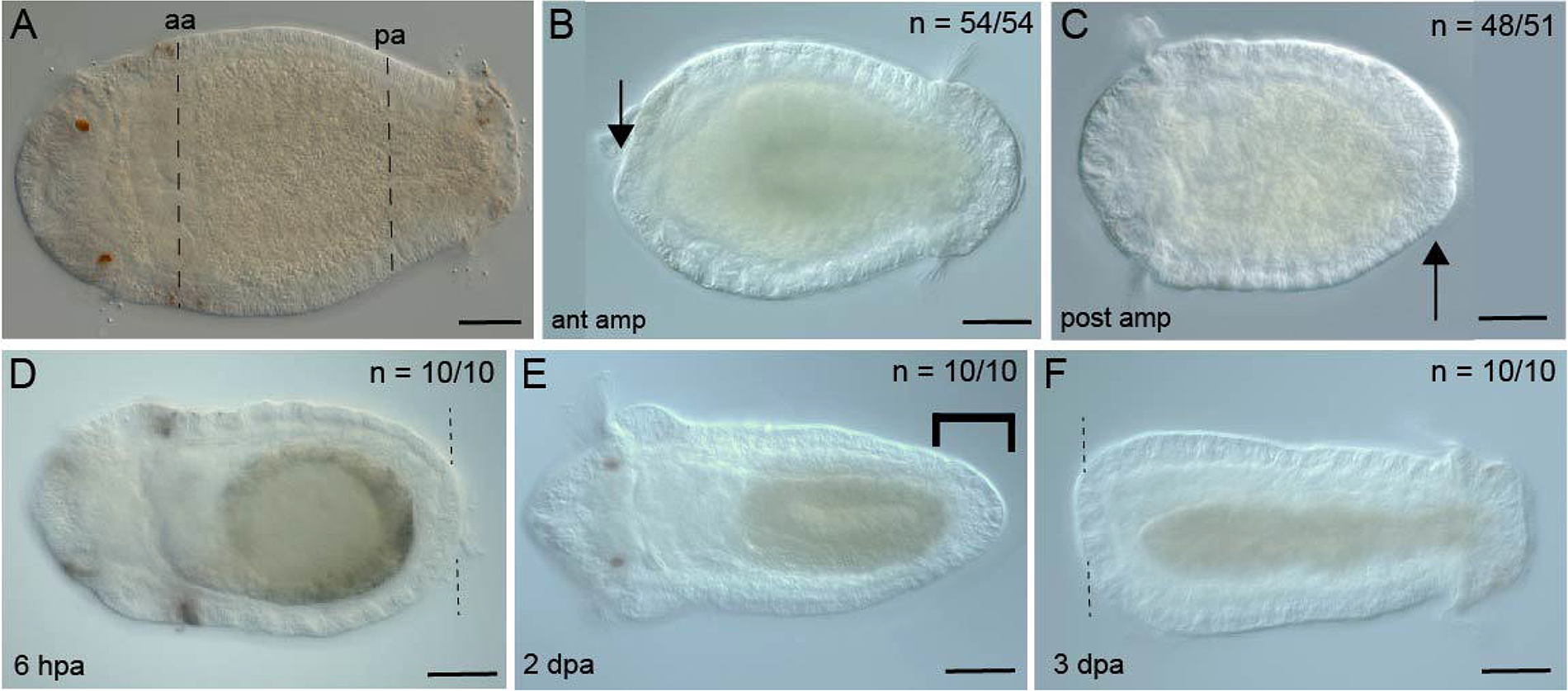
Amputated larvae undergo successful wound healing. All images are oriented in ventral view with anterior to the left. Arrows indicate the continuous layer of epithelium, indicative of successful wound healing. **A** Stage 6 uncut larva showing experiment design for amputations. The dotted lines indicate cut sites: (aa) anterior amputation and (pa) posterior amputation. **B** Stage 6 larva 6 hpa anterior amputation. **C** Stage 6 larva 6 hpa posterior amputation. **D** Stage 9 larva 6 hpa posterior amputation. **E** Stage 9 larva 2 dpa anterior amputation. **F** Larva 3 dpa anterior amputation. The black bracket represents the tissue between the gut and ectoderm. Amputation site indicated by the black dotted line. N denotes the number of animals scored that resembled the image in the panel. Scale bar is 50 µm.

Larvae survived multiple days following amputation and continued to develop towards metamorphic competency. Amputated larvae generally had high survivorship (∼75-85%) at 5 dpa, with anteriorly amputated animals and posteriorly amputated animals surviving up to 5 dpa and 7 dpa, respectively. Furthermore, amputated larvae continued to develop on schedule relative to uncut animals (Fig 2E, 2F). For example, in posteriorly amputated larvae, chaetae were present in the original tissue by 1 dpa (comparable to Stage 7), pigmentation appeared in the midgut by 2 dpa (comparable to Stage 8), and the midgut became coiled by 3 dpa (comparable to Stage 9). Anteriorly amputated larvae also developed chaetae and a pigmented midgut, but the curvature of midgut—a hallmark of Stage 9 larvae (Boyle & Seaver, 2008; Seaver et al., 2005) was delayed by one day. The growth of anterior and posterior amputations differed. The shape of the body at the cut site in anterior amputations was frequently observed to be blunt, similar in shape to a specimen following wound healing (6 hpa) (Fig 2F), whereas the tissue distal to the cut site in posterior amputations had a rounded tapered shape (Fig 2E). The tapered shape in posterior amputations is different from the shape of the wound site in aged-matched individuals (Stage 9) fixed at 6 hpa (compare Fig 2D, E). Our observations demonstrate that larvae not only survive removal of a third of their body, but they continue to develop with a similar time course to uncut animals.

### 3.4 Cell proliferation

One key feature of regeneration is the birth of new cells in the region of the wound site. Larval development is a dynamic growth process that also exhibits cell proliferation. Stage 6 and 7 larvae have a characteristic pattern of EdU incorporation following a one hour exposure to EdU. Specifically, there are numerous EdU+ cells distributed throughout the trunk, straddling the midline (Fig 3A, B). In contrast, by Stage 9, EdU+ cells are not ordinarily observed, apart from a few cells in the midgut or hindgut and posterior growth zone (PGZ) (Fig 3C, 3D). To delineate between growth and a response to amputation, we compared age-matched, uncut larvae with amputated larvae.

**Figure 3.**
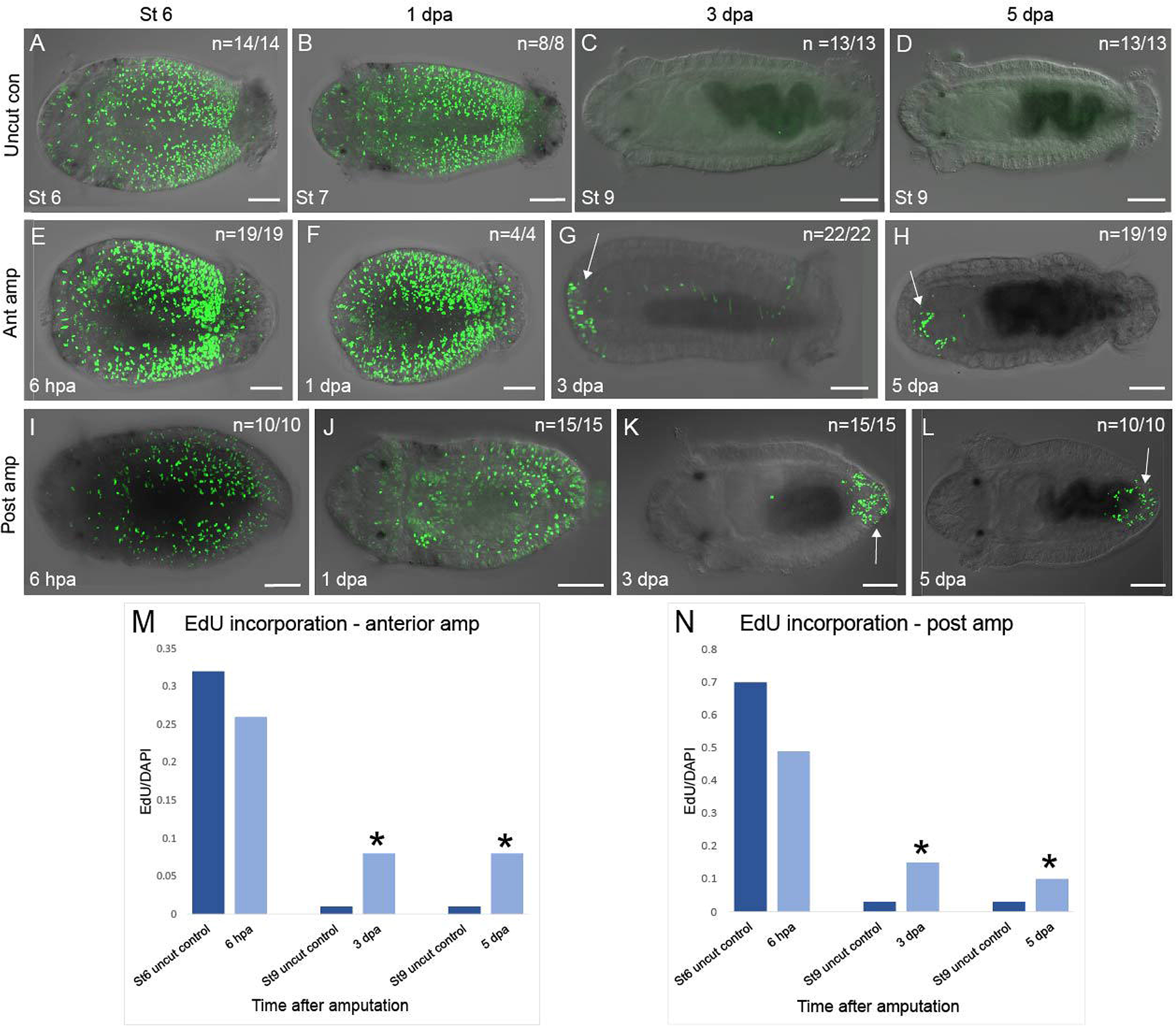
Localized cell proliferation near the amputation site in larvae. All images are oriented in ventral view with anterior to the left. Animals were exposed to EdU to label dividing cells. EdU+ cells are green. **A–D** are uncut controls: **A** Stage 6, **B** Stage 7, **C** Stage 9, and **D** Stage 9 larvae. **E–H** are anterior amputations: **E** 6 hpa, **F** 1 dpa, **G** 3 dpa, and **H** 5 dpa larvae. **I–L** are posterior amputations: **I** 6 hpa, **J** 1 dpa, **K** 3 dpa, and **L** 5 dpa larvae. **M**, **N** Quantification of EdU incorporation in anterior (M) and posterior (N) amputations and comparison with uncut controls. Time after amputation is on the X-axis and the ratio of EdU/DAPI cells is on the Y-axis. The white arrows indicate EdU+ cells near the wound site. The developmental stage or time post-amputation is indicated in the bottom left corner of each panel. N denotes the number of animals scored that resemble the image shown. Dark blue bars in the graphs represent uncut controls and light blue bars represent amputated animals. Asterisk denotes significance with p < 0.05. Ant amp, anterior amputation; Post amp, posterior amputation. Scale bar is 50 µm.

Anterior amputations show EdU incorporation near the amputation site. The initial pattern of EdU incorporation in anterior amputations (6 hpa) is similar to the pattern observed in age-matched, uncut controls (Fig 3E, 3I). This pattern of EdU incorporation persists in 1 dpa animals and in age matched controls (Fig 3B, F, J). However, by 3 dpa, EdU+ cells are localized near the wound site in the anterior ectoderm and occasionally in the mesoderm of the amputated animals (Fig 3G). EdU+ cells near the wound site persist through 5 dpa in anterior ectoderm and mesoderm (Fig 3H). In contrast, there is a notable lack of EdU+ cells in the corresponding body region of unamputated animals (Fig 3C, 3D). The ratio of EdU+ cells to nuclei is higher in amputated animals at both 3 dpa and 5 dpa at the respective amputation sites compared with the equivalent body region in uncut controls (Table 1). There is a significant difference between the number of EdU+ cells in the anterior body region anteriorly amputated larvae at 3 dpa and 5 dpa when compared to age-matched, uncut controls (One-way ANOVA test) (Fig 3M).

**Table 1.** Mean values of EdU+ cells in anterior amputations.

| Anterior Amp |  | Anterior |  | Middle |  | Posterior |  |
| --- | --- | --- | --- | --- | --- | --- | --- |
|  |  | Uncut | Cut | Uncut | Cut | Uncut | Cut |
| 6 hpa | # EdU | 133 | 102 | 466 | 235 | 387 | 136 |
|  | # Nuclei | 418 | 392 | 1216 | 417 | 551 | 250 |
|  | % | 32% | 26% | 38% | 56% | 70% | 54% |
| 3 dpa | # EdU | 3 | 29 | 18 | 2 | 14 | 24 |
|  | # Nuclei | 447 | 363 | 856 | 457 | 413 | 185 |
|  | % | 1% | 8% | 2% | 0% | 3% | 13% |
| 5dpa | # EdU | 3 | 28 | 18 | 5 | 14 | 10 |
|  | # Nuclei | 147 | 359 | 856 | 516 | 413 | 219 |
|  | % | 2% | 8% | 2% | 1% | 3% | 5% |

Posterior amputations also show EdU incorporation localized near the wound site. In both cut and uncut larvae, the number of EdU+ cells decrease over time in the anterior and middle sections of the animal. While the initial pattern of EdU incorporation in cells at 6 hpa appears similar to that observed in Stage 6 uncut controls (Fig 3I), there is a progressive shift toward expression localized to the cut site in amputated larvae. Animals at 1 dpa have fewer EdU+ cells in the anterior 2/3 of the body (Fig 3J) in comparison with uncut Stage 7 controls. By 3 dpa, EdU incorporation is not detected through the anterior 2/3 of the body, and EdU+ cells are restricted to the area surrounding the cut site (Fig 3K). This localized pattern of EdU incorporation persists through 5 dpa (Fig 3L). Furthermore, cut larvae have more EdU+ cells in the posterior region than the uncut controls (Table 2). The number of EdU+ cells in posterior sections of 3 dpa and 5 dpa larvae is significantly higher compared to age matched, uncut controls (One-way ANOVA test) (Fig 3N). Additionally, there is a higher ratio of EdU+ cells/nuclei localized to the wound site in posterior amputated larvae relative to anterior amputations. Together, these data show a marked increase in cell proliferation at the wound site following both anterior and posterior amputations.

**Table 2.** Mean of EdU+ cells in posterior amputations.

| Posterior Amp |  | Anterior |  | Middle |  | Posterior |  |
| --- | --- | --- | --- | --- | --- | --- | --- |
|  |  | Uncut | Cut | Uncut | Cut | Uncut | Cut |
| 6 hpa | # EdU | 133 | 70 | 466 | 217 | 387 | 225 |
|  | # Nuclei | 418 | 155 | 1216 | 556 | 551 | 463 |
|  | % | 32% | 45% | 38% | 39% | 70% | 49% |
| 3 dpa | # EdU | 3 | 2 | 18 | 6 | 14 | 78 |
|  | # Nuclei | 447 | 306 | 856 | 702 | 413 | 525 |
|  | % | 1% | 1% | 2% | 1% | 3% | 15% |
| 5dpa | # EdU | 3 | 0 | 18 | 0 | 14 | 51 |
|  | # Nuclei | 147 | 331 | 856 | 547 | 413 | 513 |
|  | % | 2% | 0% | 2% | 0% | 3% | 10% |

### 3.4 Expression of the stem cell marker vasa

To further characterize the larval response to amputation, expression of the stem cell marker *vasa* was analyzed by *in situ* hybridization (Dannenberg & Seaver, 2018; Juliano et al., 2010). In uncut larvae, *vasa* expression is dynamic from Stage 6 through Stage 9 (Dill & Seaver, 2008). In Stage 6 larvae, there is expression in the foregut, the mesoderm of the trunk and the PGZ (Fig 4A). By Stage 8, expression is restricted to the foregut, the multipotent progenitor cell cluster (MPC), and the PGZ (Fig 4B). This pattern continues through Stage 9 (with some variability in detection of the MPC) (Fig 4C).

**Figure 4.**
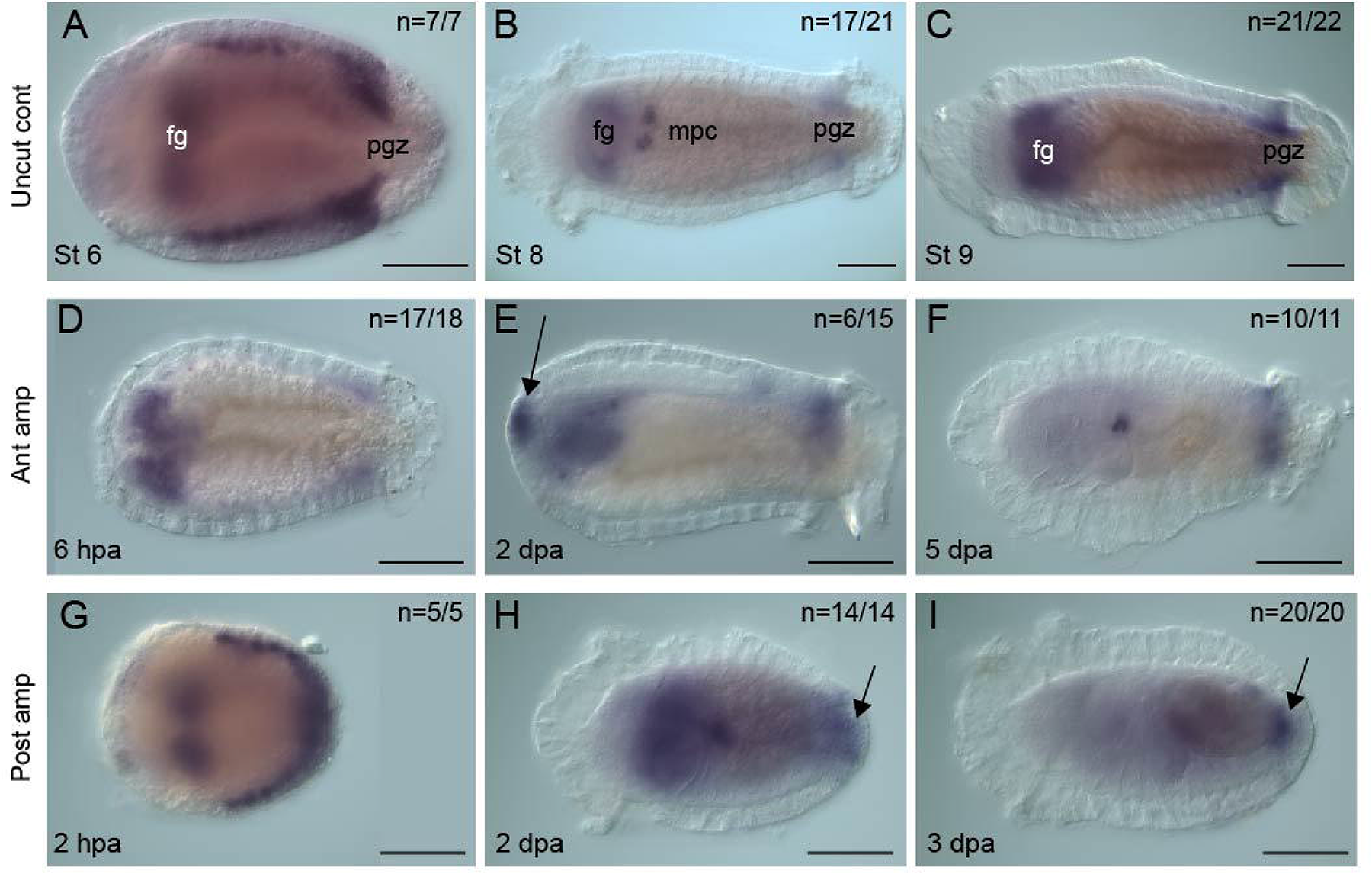
*Vasa* is expressed at the wound site in amputated larvae. All images are oriented in ventral view with anterior to the left. The purple coloration indicates localization of *vasa* transcripts by in situ hybridization. **A**, **B**, and **C** *Vasa* expression patterns in uncut larvae at Stage 6, Stage 8, and Stage 9. **D**, **E**, and **F** are anterior larval amputations. **G**, **H**, and **I** are posterior larval amputations. **D** and **G** are amputated larvae 6 hpa (D) and 2 hpa (**G)**. **E** *De novo* expression is observed in anterior amputations at 2 dpa, as indicated by the arrow and is no longer detectable by 5 dpa (**F)**. In posterior amputations, expression near the wound site is observed at both 2 dpa (**H**) and 3 dpa (**I**), as indicated by arrows. The time post-amputation is indicated in the bottom left corner of each panel in D–I. N denotes the number of animals scored that resemble the image shown. Ant amp, anterior amputation; Fg, foregut; mpc, multi-progenitor cell cluster; Post amp, posterior amputation; pgz, posterior growth zone. Scale bar is 50 μm.

*Vasa* expression is detected at the wound site in both anterior and posterior amputations. Initially, the *vasa* expression pattern in both amputation groups is very similar to the observed pattern in uncut controls (6 hr post amputation) (Fig 4D). However, by 2 dpa, there is *de novo* expression in the anterior ectoderm at the amputation site of anteriorly cut larvae (Fig 4E). By 5 dpa, expression at the wound site is no longer detected, although expression in the MPC and PGZ persists (Fig 4F). In posterior amputations at 2 hpa, trunk mesodermal expression is continuous from one side of the animal to the other and wraps around the posterior end of the body (Fig 4G). We interpret this expression along the posterior face of the animal to be the result of wound healing that pulls the two lateral trunk domains of expression together. At 2 dpa, *vasa* is expressed in the posterior mesoderm and ectoderm near the wound site (Fig 4H). This expression near the wound site is also detectable 3 dpa (Fig 4I) and 4 dpa (data not shown). In summary, *vasa* expression is detected at the wound site 2–3 dpa for both anterior and posterior amputations.

### 3.5 Generation of new tissue

The presence of localized cell proliferation at the wound site (Fig 3G, 3H, 3K, and 3L) and what appears to be growth of new tissue (Fig 2E) warranted a more detailed investigation of the tissue surrounding the wound site. To distinguish between pre-exiting tissue and new tissue, the location of the cut site was determined using the nervous system and musculature markers. We used a framework commonly used for identifying the cut site in juvenile worms with an anti-acetylated tubulin antibody marking the nervous system (Fig 1B). An abrupt change in the thickness of the connective nerves was used to indicate the position of the cute site (Fig 1D, 5B, 5B’, and 5C). An abrupt decrease in the density of muscle fibers also indicated the location of the cut site (Fig 1E, Fig 5F and 5F’). Typically, there was a precise concordance in position between an abrupt change in muscle fiber density and thickness of connective nerves. Therefore, we could consistently identify the location of the cut site using these two markers.

**Figure 5.**
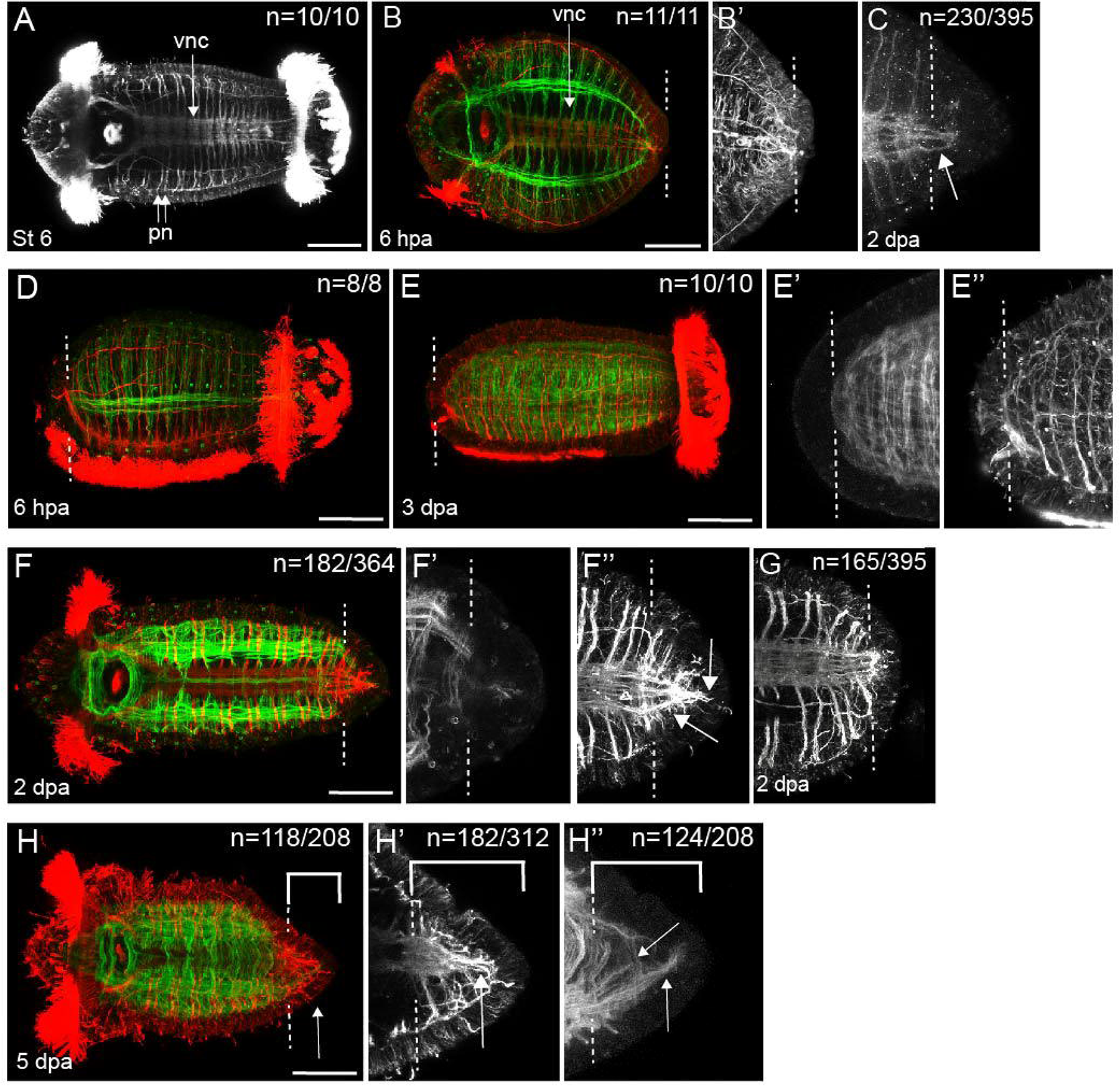
Tissue response to amputation. **A–C** and **F–I”** are oriented in ventral view. **C–D”** are oriented in lateral view. In all images, anterior is to the left. **A** uncut Stage 6 larva labeled with anti-acetylated tubulin to visualize the nervous system and ciliary bands. Larvae are labeled with anti-acetylated tubulin (red) and phalloidin (green) showing the nervous system and musculature for all panels except A, C and G. Larvae shown in panels A, C and G were labeled with anti-acetylated tubulin. **B** and **B’** are the same specimen 6 hpa. **C** neural fibers are present distal to the cut site in 2 dpa posterior amputee (arrow). **D** larva 6 hpa after anterior amputation. **E**, **E’** and **E”** are the same specimen, 3 dpa anterior amputation. **E’** magnified view of muscle fibers as marked by phalloidin. **E”** magnified view of anti-acetylated tubulin labeling. **F**, **F’**, and **F’’** are the same larva 2 dpa. **F** Nervous system and musculature in a larva 2 dpa. **F’** magnified view of posterior wound site showing few muscle fibers distal to cut site. **F”** magnified view of posterior wound site showing fibers in the tissue distal to the amputation site (arrows). **G** shows a specimen lacking neural fibers distal to the cut site. **H**, **H’**, and **H”** are the same individual at 5 dpa posterior amputation. **H’** neurites distal to the cut site (arrow). **H’’** longitudinal and circular muscles are present distal to the cut site (bracket). White brackets in H, **H’** and **H”** indicate tissue distal to the amputation site. N denotes the number of animals scored that resemble the representative image. Amputation site is indicated by a white dotted line. vnc, ventral nerve cord; pn, peripheral nerves. Scale bar is 50 µm.

The identification of the cut site in amputated larvae allowed us to identify new tissue and quantify tissue distal to the amputation site. No evidence of significant growth of tissue was detected in anterior amputations between 6 hpa and 3 dpa (see Table 3). In contrast, there was significant growth in posterior amputations between 6 hpa and 5 dpa. Additional measurements of posterior amputations indicated a significant change in body width (body width at the mouth compared to immediately posterior to the cut site) between 6 hpa and 5 dpa. This information is consistent with the presence of new growth in posterior amputations; regenerated tissue is often more narrow than the width of the original tissue. While the position of the cut site indicated by muscle and nerves indicated new growth, there was one measurement that did not. Measurements from the posterior end of the pigmented gut to the posterior of the animal for both the 6 hpa stage 9 and 5 dpa larvae did not show a significant difference between these two groups.

**Table 3.** Measurement of tissue distal to cut site.

|  |  | **Muscle** | **Nervous System** | **Gut** | **Width 2 /Width 1** |
| --- | --- | --- | --- | --- | --- |
| **Ant Amp** | **6 hpa** | 19 $\mu\text{m}$ | 18 $\mu\text{m}$ | N/A | N/A |
| | **3 dpa** | 22 $\mu\text{m}$ | 15 $\mu\text{m}$ | N/A | N/A |
| **Post Amp** | **6 hpa** | * 17 $\mu\text{m}$ | * 12 $\mu\text{m}$ | N/A | 0.675 |
| | **5 dpa** | * 62 $\mu\text{m}$ | * 67 $\mu\text{m}$ | * 45 $\mu\text{m}$ | 0.453 |
| | **St 9 6 hpa** | N/A | N/A | * 37 $\mu\text{m}$ | N/A |

\* = significance, $p < .05$

We next characterized the nervous system and musculature of the tissue distal to the cut site. Similar to what was observed in 6 hpa control larvae (Fig 5D), anterior amputations in 3 dpa larvae lacked clear evidence of neurites and muscle fibers distal to the wound site (Fig 5 E–E”). In posteriorly amputated larvae, a mixture of results was observed. In 2 dpa posteriorly amputated larvae with a discernable cut site and new growth, neuronal projections distal to the amputation site were observed that were not detected in 6 hpa larvae (Fig 5B, 5F, 5F’, 5F’’). There were also cases in which there was seemingly no change in the larval nervous system (Fig 5G; n = 165/395). Cases in which new growth was not observed lend confidence to those instances when we observed processes distal to the cut site (compare Fig 5F’’ and 5G). At 5 dpa, nerve processes persisted in tissue distal to the cut site (Fig 5H, 5H’). These longitudinal nerve processes are located at the midline and span the length of the new tissue. Additionally, there were circumferential processes similar to the orientation of peripheral nerves, although these processes were somewhat disorganized relative to peripheral nerves in uncut animals (Fig 5H’). These neuronal processes are oriented medial to lateral, extending from the midline. The entire region of the new tissue has a lower density of muscle fibers relative to the preexisting tissue (Fig 5B, 5H’). The muscle fibers that are present are predominately longitudinal muscle fibers. In many cases, individual longitudinal muscle fibers appear to extend from preexisting tissue into tissue distal to the amputation site (Fig 5H”, arrow), whereas circular muscle fibers were only occasionally observed posterior to the amputation site. These circular fibers are unlikely associated with a telotroch because amputated larvae lack evidence of regrown ciliary bands. A similar arrangement of muscle and nerve fibers in the tissue distal to the amputation site is present in animals 7 dpa (data not shown). While we observed multiple tissue types distal to the cut site, we did not observe evidence of new segmentation. Conventionally, newly formed segments are characterized by the appearance of new ganglia, regularly spaced circumferential peripheral nerves and segmental boundaries. The nervous system distal to the cut site did not organize into ganglia by 7 dpa, and peripheral nerves were disorganized. We did not observe segmental boundaries in the ectoderm. In conclusion, new tissue growth was observed along with changes in organization of the nervous system and musculature in posteriorly amputated larvae, although there was a lack of new tissue in anteriorly amputated larvae.

### 3.6 Neural specification and differentiation

Next, we sought to determine if the neural and muscle cell types observed distal to the amputation site resulted from the birth of new cells. We rationalized that expression of molecular markers of cell fate specification and differentiation in the tissue distal to the amputation site indicated birth of new cells. For the nervous system, gene expression patterns of neural specification (*ash1*) and neural differentiation markers (*elav*) were analyzed near the wound site (Meyer & Seaver, 2009). In *C. teleta*, *ash1* is a marker of neural specification during initial neurogenesis of the brain and VNC (Meyer & Seaver, 2009; Sur et al., 2017). In Stage 6 larvae, *ash1* is expressed in the brain, foregut, ventral neural ectoderm, PGZ, and the mesoderm in the pygidium (Fig 6A). By Stage 9, *ash1* expression is restricted to the foregut and PGZ (Fig 6B). In anterior amputations, the pattern observed at 6 hpa for *ash1* expression mirrors that of uncut Stage 6 larvae, with the notable exception of missing brain expression due to the removal of the head (Fig 6C). In 3 dpa specimens, *ash1* is expressed in both the anterior ectoderm and pharynx (Fig 6D). Expression in the ectoderm can be regarded as *de novo ash1* anterior expression since age-matched, uncut controls lack anterior ectoderm expression (compare panels Fig 6B and 6D, E) (n =32/39). The novel anterior ectodermal expression of *ash1* at 3 dpa was present in a punctate pattern similar to the pattern observed in the anterior ectoderm during *C. teleta* brain development. In contrast, expression in the pharynx corresponds with expression in age matched uncut controls, albeit with fewer cells.

**Figure 6.**
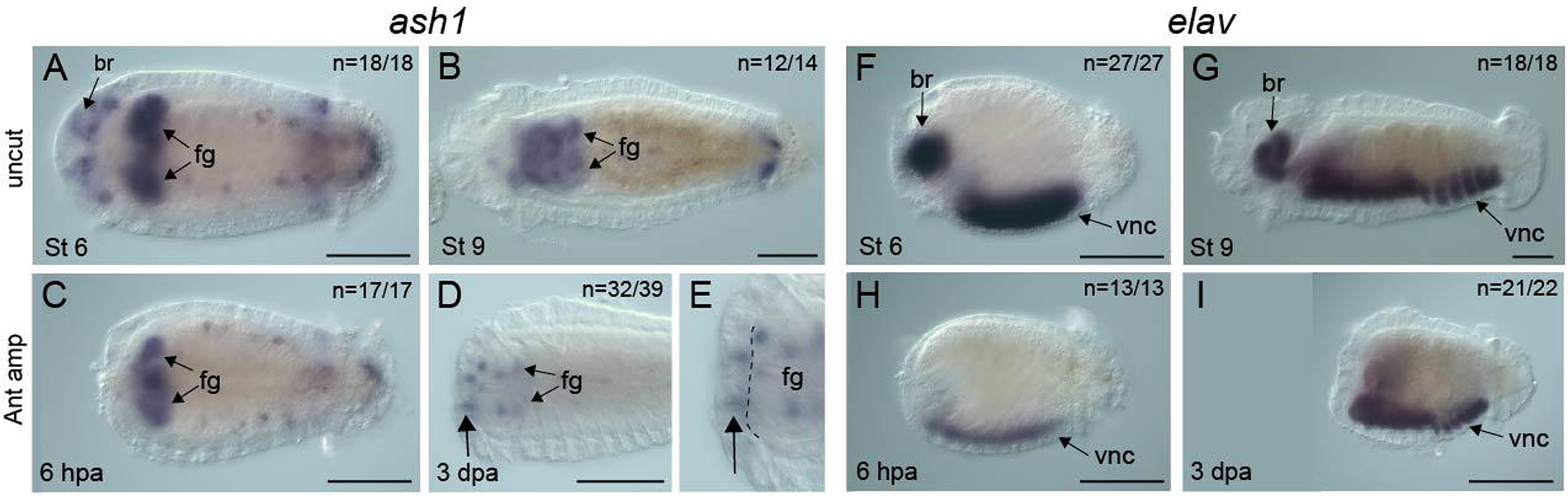
Amputation induces *ash1* but not *elav* expression at the anterior wound site. All images are oriented with anterior to the left. **A-E** are ventral views and **F-I** are lateral views. The purple coloration indicates expression of either *ash1* (**A-E**) or *elav* (**F-I**) transcripts. **C-E** and **H-I** are anterior amputations. **A** Stage 6 uncut control. **B** Stage 9 uncut control. **C** 6 hpa. **D** and **E** are the same 3 dpa individual. Vertical arrows indicate novel *ash1* ectodermal expression. The boundary between ectodermal and mesodermal layers is indicated by a black dotted line. **F** Stage 6 and **G** Stage 9 uncut controls. **H** 6 hpa. **I** 3 dpa. N denotes the number of animals scored that resemble the image indicated. Ant amp, anterior amputation; br, brain; fg, foregut; vnc, ventral nerve cord. Scale bar is 50 µm.

After observing novel anterior neural specification following amputation, the next step of neurogenesis was analyzed by characterizing the expression pattern of *elav*, a marker of differentiating neurons. Between Stages 6 and 9 in uncut controls, *elav* is expressed in the brain and the VNC (Fig 6F & G). In 6 hpa anterior amputations, *elav* is expressed only in the VNC (Fig 6H). This expression pattern persists through 3 dpa, with no new expression domains observed in amputated animals (n =21/22) (Fig 6I). In the one differing case of the 22 scored animals, there were a few *elav*-expressing cells in the anterior ectodermanteriorly located cells. These cells are part of the pharynx that abnormally everted during morphogenesis.

### 3.7 Amputation induces expansion of MyoD expression

We also examined the expression of *MyoD*, a gene known for its role in muscle specification and differentiation (Zammit, 2017). In uncut larvae, *MyoD* is expressed in the mesoderm of the posterior trunk segments at Stage 6 (Fig 7A) and Stage 9 (Fig 7B). There is a developmental progression from anterior to posterior in the larval trunk, with the more mature segments having an anterior position (Seaver et al., 2005). Thus, *MyoD* is expressed in the younger and newly forming segments. Six hours after both anterior and posterior amputations (6 hpa), expression is restricted to the posterior portion of the trunk (Fig 7 C & E), similar to what is observed in uncut controls. At 7 dpa in anterior amputees (Fig 7D) and at 3 dpa in posterior amputees (Fig 7F), there is a drastic expansion of *MyoD* mesodermal expression along the entire length of the trunk. This expansion in *MyoD* expression is in pre-existing tissue in older segments and includes tissue distant from the wound site in the case of posterior amputations.

**Figure 7.**
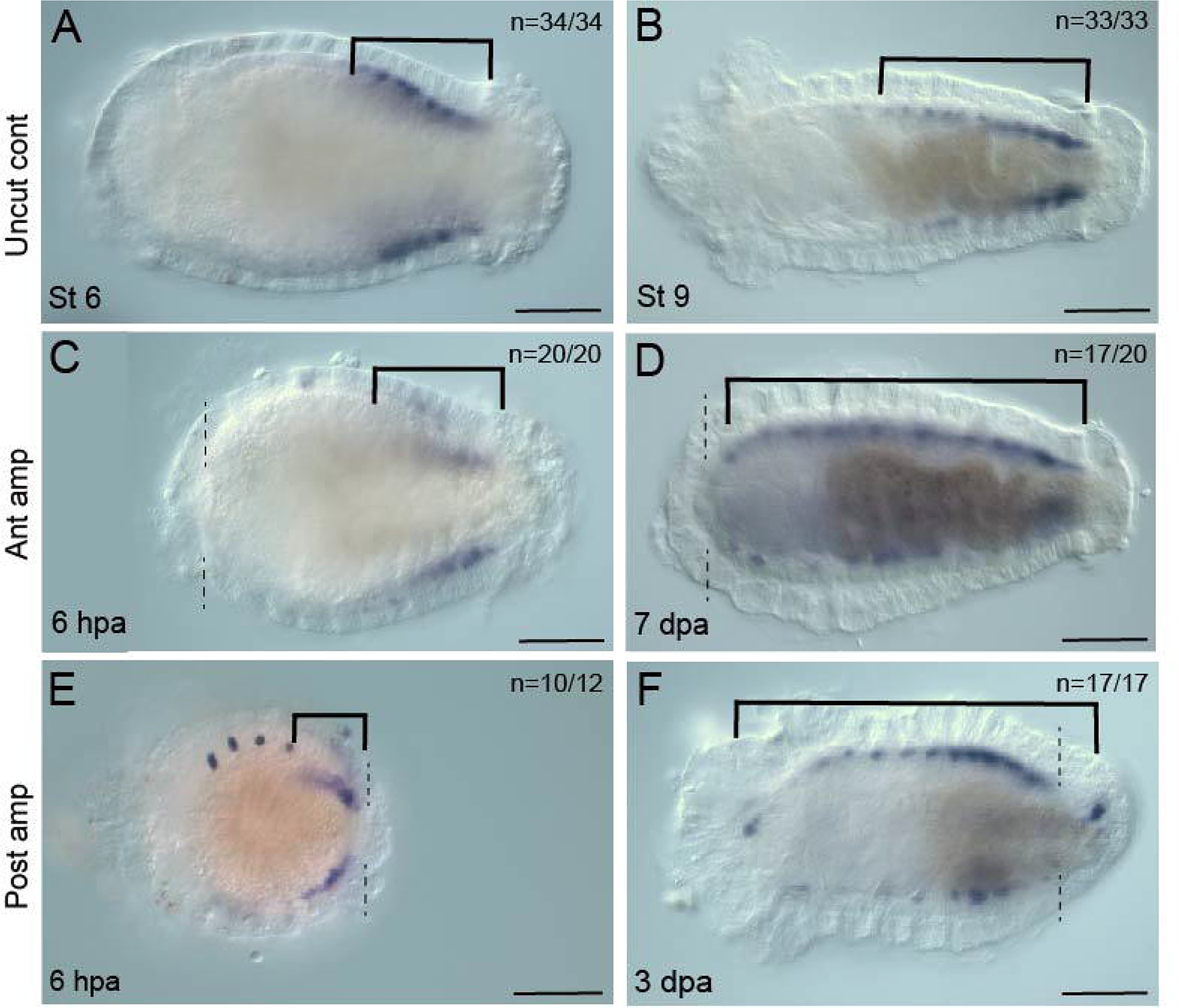
Amputation induces expansion of *MyoD* expression. All images are oriented in ventral view with anterior to the left. Localization of *MyoD* transcripts is visualized by the purple coloration. **A** Stage 6 and **B** Stage 9 uncut controls. *MyoD* is expressed in the posterior trunk mesoderm. **C** and **D** are anterior amputations (ant amp). **C** 6 hpa. **D** anterior shift of anterior boundary of *MyoD* expression is observed at 7 dpa. **E** 6 hpa posterior amputation (post amp). Spots of staining anterior to the bracket is nonspecific trapping in the chaetae. **F** *MyoD* is expressed along the length of the body in 3 dpa animals (compare with E). Brackets indicate the regions of detectable *MyoD* expression along the anterior-posterior axis. Amputation sites are indicated by black dotted lines. N denotes the number of animals scored that resemble the representative image. Ant amp, anterior amputation; Post amp, posterior amputation. Scale bar is 50 μm.

### 3.8 Rare regeneration of differentiated cell types

Following amputations, a few differentiated cell types occasionally regenerated, namely the larval eyes and cilia. Larvae have two cerebral eyes, with each eye consisting of three cells: a pigment cell (Fig 8A), a sensory cell (Fig 8C), and a support cell (Yamaguchi & Seaver, 2013). The pigment cell is readily visible by its orange pigment granules and the sensory cell can be visualized by immunolabeling with the antibody 22C10 (Yamaguchi & Seaver, 2013). Currently, we are unable to assess the presence of the support cell. In uncut Stage 9 larvae, the pigment cells are located slightly anterior, if not inline, with the prototroch, in a lateral position (Fig 8A). The sensory cell in uncut Stage 9 larvae (Fig 8C) is located immediately adjacent and lateral to the pigment cell and has an axonal projection extending from the cell body to the external surface of the ectodermal epithelium. The pigment and sensory cells were scored as independent characters. Following head amputation, a few pigment cells regenerated, and they are variable in location and number (n = 28/420; 3-5 dpa). Additionally, the pigment cells in amputated larvae are ectopically positioned. For example, as seen in Fig 8B, there is a pigment cell in proximity to the midgut. Regenerated sensory cells were also observed in amputated larvae and also exhibited variability in location (n= 8/220, 3 –7 dpa). Axon projections in regenerated sensory cells extended to the outer ectodermal surface of the animals (Fig 8D). There was a higher percentage of cases with regenerated pigment cells (7%) compared with sensory cells (4%) in anterior amputations (Fig 8G). Summation of the total number of pigment cells and sensory cells indicates that at least one cell of the eye regenerated in 6% of cases amputated (n = 36/640).

**Figure 8.**
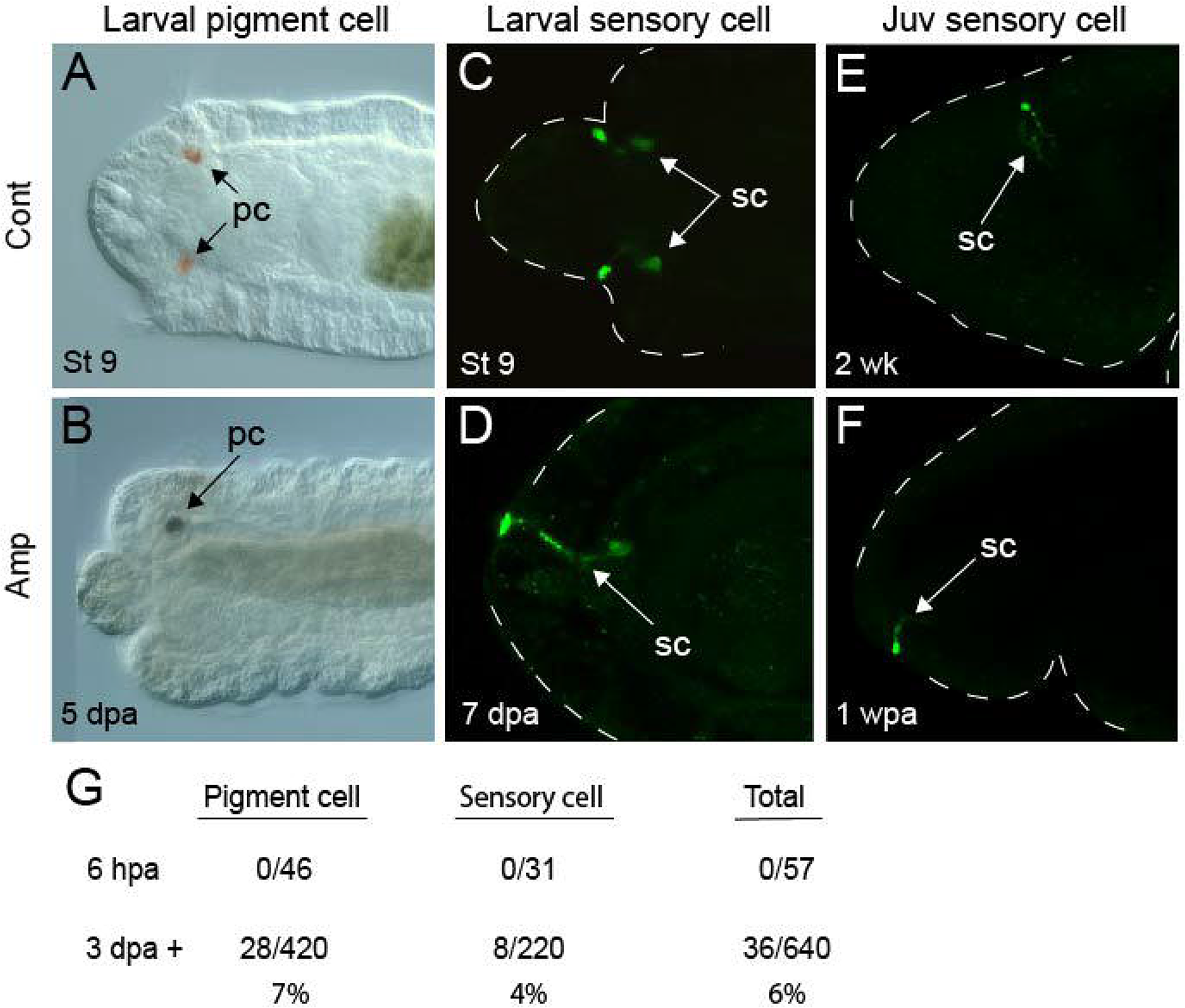
Rare eye reformation following anterior amputation. All images are oriented with anterior to the left. **A** – **D** are ventral views. **A** DIC image of the anterior end of a Stage 9 uncut larva. **B** DIC image of the anterior end of a larva 5 days after anterior amputation. The image shown is a merge of multiple focal planes. The pigmented cells (pc) of the eye are indicated by arrows. **C**–**F** eye sensory cells (sc, white arrows), labeled by the 22C10 antibody (green). **C** Uncut Stage 9 control. **D** anterior amputated larva 5 dpa. **E** and **F** are juvenile worms in lateral view. **E** uncut 2 week-old juvenile and **F** juvenile 1 week post-amputation of the head. **G** Table displaying the number of pigment cells, sensory cells, or either cell (total) present in anteriorly amputated larvae at 6 hpa and 3 dpa or later. The dotted line indicates the outline of the animal. Juv, juvenile; mg, midgut; pc, pigment cells; sc, sensory cells.

We were surprised by the appearance of new larval eye cells in amputated *C. teleta* larvae since regeneration of anterior structures is not reported in adults (Yamaguchi & Seaver, 2013). Therefore, we conducted a similar experiment with juveniles. Four-week-old juveniles were amputated immediately anterior to the mouth, removing the eye cells. After screening for successful wound healing and visual inspection that pigment cells were removed, a subset of the amputated juveniles was fixed at 4 hpa to confirm the removal of the sensory cells (n =21/21). No pigment cells were observed in amputated juveniles at 7 dpa (n = 0/43). However, a small proportion of cases with eye sensory cells were observed (Fig 8E) (n = 6/43; 14%). Regenerated sensory cells were ectopically located (compare Fig 8E and 8F). Five of the six juveniles with regenerated sensory cells had a small portion of brain present in the anterior of the animal; only one case lacking a brain had a sensory cell. In summary, pigment cells did not regenerate in juveniles, although sensory cells were detected in a small fraction of cases. These rare instances of eye sensory cell regeneration represented the most dramatic phenotype observed following juvenile anterior amputations.

There are four external ciliated structures in *C. teleta* larvae: prototroch, neurotroch, telotroch, and pygidial cilia (Fig 9A, B). The prototroch and telotroch are composed of relatively long cilia (prototroch cilia length ∼25-30 microns and telotroch cilia length ∼20-28 microns), and the neurotroch and pygidium have short cilia (neurotroch cilia length ∼ 8-10 microns, pygidium length ∼5-7 microns). At 6 hpa, larvae lacked long cilia at the cut site (Fig 9 C and 9D), due to amputation of the prototroch and telotroch in anterior and posterior amputations, respectively. At 5 dpa, short cilia (approximately 8 microns in length) were observed in rare cases (n = 12/161; 7%) at the anterior end of anteriorly cut animals, distal to the amputation site (compare Fig 9E and F). Long cilia were not observed following anterior amputations (n = 0/161). While rare (n = 22/370; 6%), both long (∼25 microns) and short (∼9 microns) cilia appeared distal to the cut site 3–7 days after posterior amputations (compare Fig 9G with 9H and 9H’).

**Figure 9.**
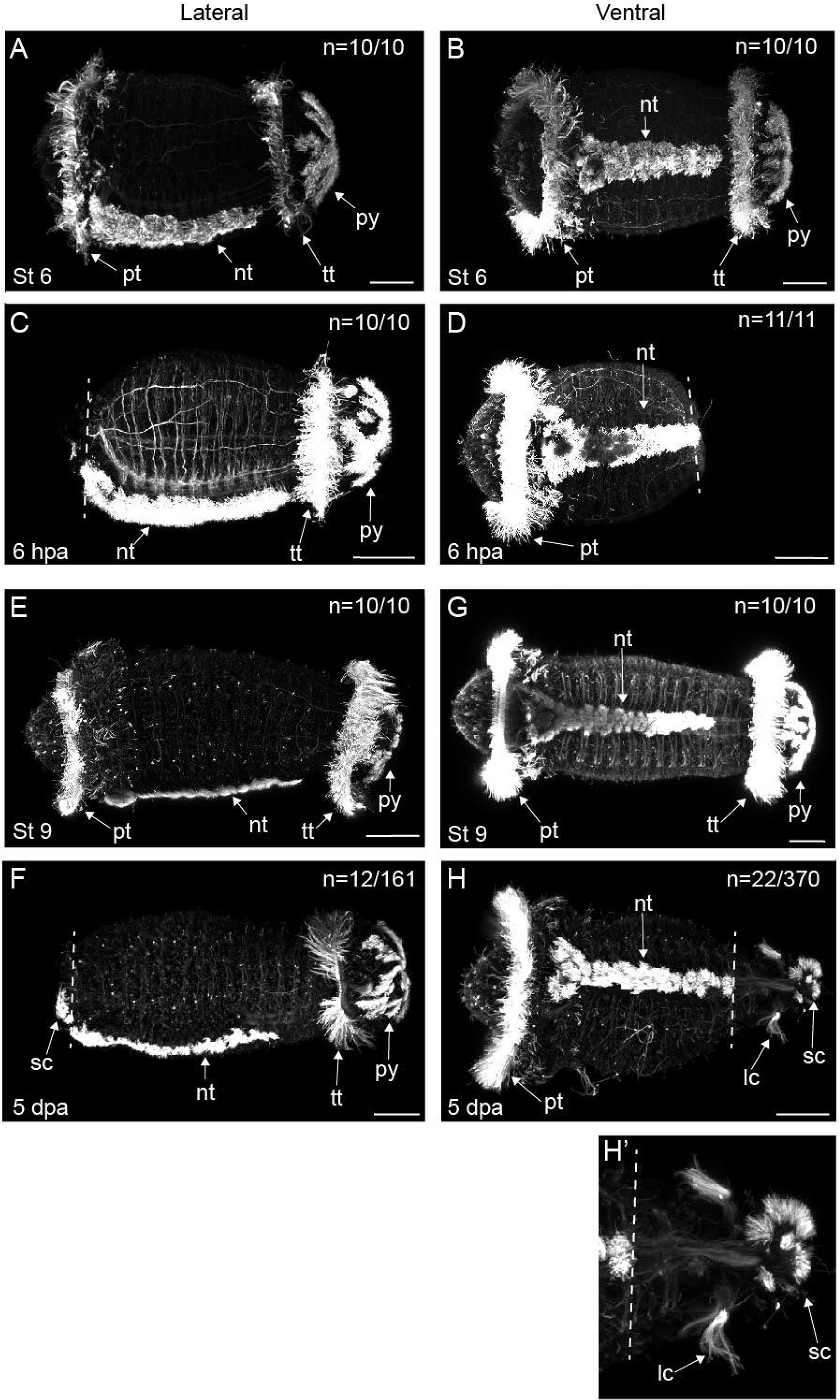
Rare reformation of cilia occurs in amputated larvae. Animals are labelled with anti-acetylated tubulin (white). All images are oriented in with anterior to the left. **A, C, E, B,** and **F** are oriented in lateral view. **B, D, G, H,** and **H’** are oriented in ventral view. **A** Stage 6 uncut larva. **D** Stage 9 uncut larva. **B** and **E** are anterior amputations. **B** larva 6 hpa. **E** larva 5 dpa with short cilia visible distal to the cut site (arrow). **D, H,** and **H’** are posterior amputations in ventral view. **D** larva 6 hpa. **H** and **H’** larva 5 dpa with both long cilia (LC) and short cilia (SC) distal to the cut site (arrows). **H’** is a magnification of **H**. N denotes the number of animals scored that are similar to the image represented. Amputation site is indicated by the white dotted line. Ant amp, anterior amputation; lc, long cilia; nt, neurotroch; post amp, posterior amputation; pt, prototroch; pyg, pygidium; sc, short cilia; tt, telotroch. Scale bar is 50 µm.

### 3.9 Metamorphosis following amputation

*C. teleta* is competent and can be induced to undergo metamorphosis into juveniles as Stage 9 larvae (Burns et al., 2014). Following amputation, larvae were screened for stereotypical signs of successful settlement and metamorphosis following the definition of Cohen and Pechenik (1999). These changes include a loss of swimming and initiation of burrowing behavior, loss of both ciliary bands (telotroch and prototroch), and the appearance of an elongated, worm-like body shape. Uncut, uninduced control larvae swam with their prototroch and telotroch bands intact for the duration of the experiment (Fig 10A) (2 hr, n = 532/534). In contrast, virtually all larvae in the induced, unamputated control group successfully underwent metamorphosis within two hours of exposure to the metamorphic cue, settling on the bottom of the dish, dropping their cilia, adopting burrowing behavior, and extending in length along the anterior-posterior axis (Fig 10B) (n = 203/207). Both sets of amputated larvae followed a similar pattern to what was observed for unamputated larvae. Specifically, one day after anterior amputation, larvae either continued to swim in the water column in the absence of a metamorphic cue (Fig. 10C) (n=65/66) or settled to the bottom of the dish and underwent metamorphosis within two hours of exposure to a metamorphic cue (Fig. 10D) (n=135/136). Similarly, individuals whose posterior ends had been removed retained their prototrochal bands and continued to swim in the absence of a metamorphic cue (Fig. 10E, arrow) (n=65/65), yet underwent rapid settlement and metamorphosis within two hours of exposure (Fig. 10F) (n=143/144). In the absence of an induction cue, amputated larvae continued to swim in the water column and retained their ciliary bands for several days, demonstrating that amputation itself does not induce metamorphosis (Fig 9E and 9F). Together, these results demonstrate *C. teleta* larvae can detect a metamorphic cue and successfully execute metamorphosis in the absence of either anterior or posterior structures.

**Figure 10.**
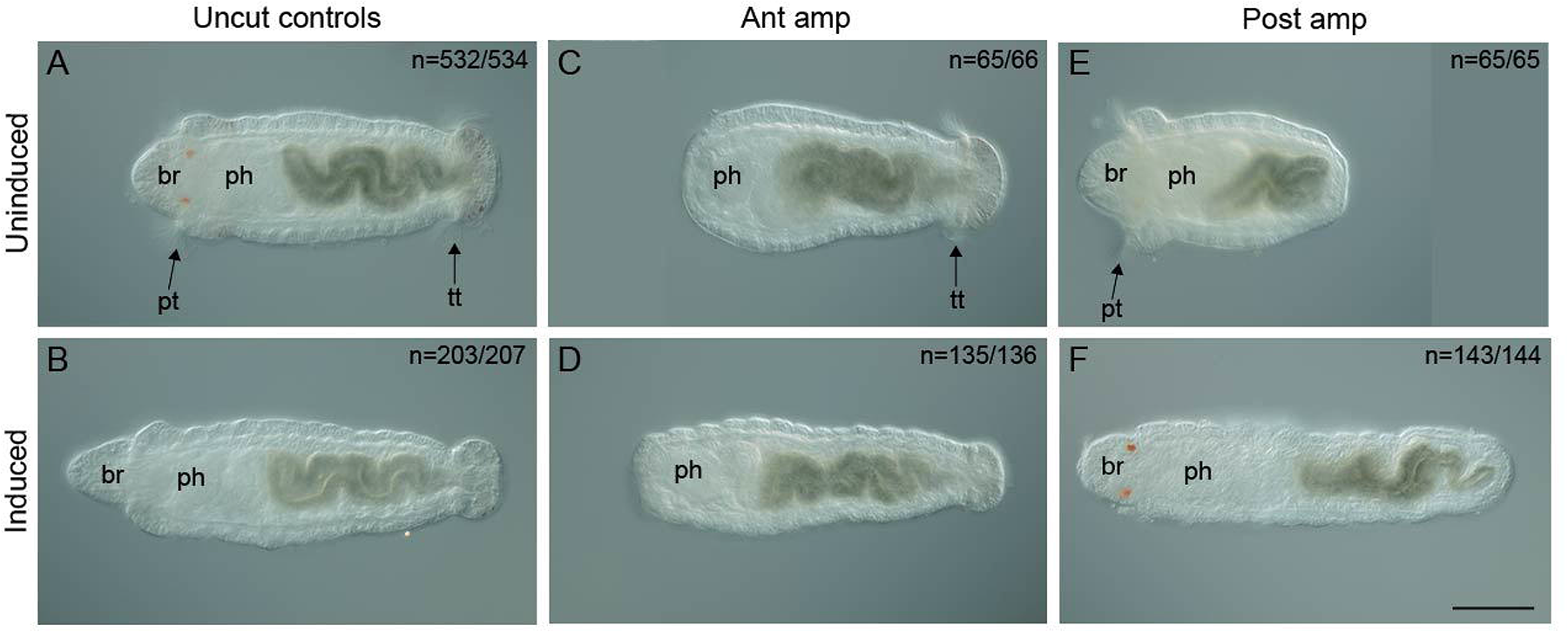
Induction of metamorphosis in amputated larvae. All images are oriented in ventral view with anterior to the left. **A** Uncut Stage 9 larva. **B** Uncut animal 2 hr after induction with metamorphic cue. **C** anteriorly amputated larva not exposed to metamorphic cue. **D** anteriorly amputated animal 2 hr after exposure to metamorphic cue. **E** posteriorly amputated larva not exposed to metamorphic cue. **F** posteriorly amputated animal 2 hr after exposure to metamorphic cue. N denotes the number of animals scored that are similar to the image shown. Ant amp, anterior amputation; Br, brain; ph, pharynx; post amp, posterior amputation; pt, prototroch; tt, telotroch. All images have the same magnification; scale bar, 50 µm.

## 4. Discussion

### 4.1 Limited regeneration in C. teleta larvae

This study demonstrates that the initial events in response to amputation appear to be quite similar between *C. teleta* larvae and juveniles. Following both anterior and posterior amputation, larvae wound heal in a similar time frame as previously reported for posterior amputation of juveniles (i.e., within 6 hpa) (de Jong & Seaver, 2016). Following wound healing, EdU+ cells appear localized at the wound site at 2 dpa in juvenile posterior regeneration, and the number of EdU+ cells increase over time (de Jong & Seaver, 2016). EdU incorporation in anterior-facing wound sites in juveniles of *C. teleta* has not been formally characterized. In both anterior and posterior larval amputations, EdU is incorporated near the wound site by 3 dpa and this persists through 5 dpa, comparable to the pattern of EdU incorporation at the wound site in regenerating juveniles. There is a difference in the number of EdU+ cells between anterior and posterior amputations in larvae, with posterior amputations exhibiting more EdU+ cells. One difference in cell proliferation patterns between juveniles and larvae is a change in position of the localized EdU+ cells. During posterior regeneration in juveniles, localization of EdU+ cells progressively shifts anteriorly to a subterminal position, a shift that corresponds with the re-establishment of the juvenile PGZ. We did not observe such a shift in larvae. It is interesting to note that in adults of the annelid *Paranais litoralis*, BrdU is incorporated near the anterior wound site even though these animals have lost the ability to regenerate their heads (Bely & Sikes, 2010). This finding, along with our results in *C. teleta* larvae, emphasize that a block in regeneration can occur subsequent to the initiation of cell division at the wound site, and cell division at the wound site is not a reliable predictor of successful regeneration.

Interestingly, expression of the stem cell marker *vasa* at the wound site was observed in both anteriorly and posteriorly amputated larvae. In successful posteriorly regenerating juveniles, *vasa* is expressed locally at the wound site at 2 dpa and persists for several days (de Jong & Seaver, 2017). In larvae, the duration of *vasa* expression and proportion of cases is different between anterior and posterior amputations. In anterior amputations, *vasa* is expressed in the anterior ectoderm in approximately 1/3 of cases and is transient, detectable at 2 dpa but not 5 dpa. Conversely, localized *vasa* expression is observed at 2 dpa and persists through 5 dpa in posterior amputations. We interpret the localized expression of *vasa* at the posterior wound site as either the result of *de novo* expression or expression maintained from an earlier stage (i.e., 6 hpa). The spatial and temporal patterns of EdU incorporation and *vasa* expression are very similar in posterior amputations. In contrast, *vasa* expression has a shorter duration than EdU incorporation near the wound site in anterior amputations. We hypothesize that these differences between anterior and posterior amputations is related to the new tissue growth in posterior amputations that is absent following anterior amputations.

Additional features of juvenile posterior regeneration are the presence of neural projections into the wound site and muscle reorganization distal to the amputation site. In juveniles, this phenomenon is observed approximately at 2 dpa (de Jong & Seaver, 2016). In larval posterior amputations, neurites are present distal to the cut site by 2 dpa. However, these neurites were not observed in larval anterior amputations. With respect to muscle fiber organization, there is an abrupt transition between the presence and absence of muscle fibers distal to the cut site in both juveniles and larval posterior amputations that was not observed in larval anterior amputations. We were unable to determine whether fibers present distal to the amputation site in posterior larval amputations resulted from birth of new neural or muscle cells or from the extension of pre-existing cells.

To better understand the characteristics and development of the tissue distal to the cut site in larvae, we characterized the specification and differentiation of cell types (i.e., neural cell types and cilia) in amputated larvae. The timing of neural specification or differentiation in juveniles has not been investigated in detail, but differentiated tissues are present by 7 dpa (de Jong & Seaver, 2016). Analysis of *de novo ash1* expression indicates the specification of neural fates in larval anterior amputations. Surprisingly, subsequent differentiation of neural tissues was not detectable, as illustrated by the absence of *elav* expression in larval amputations. Anterior amputation of larvae results in incomplete tissue and structure regeneration—an observation consistent with the response in juveniles to anterior amputations. Despite a general lack of anterior structural differentiation in amputated larvae, we surprisingly observed rare cases in which differentiated eye cells and cilia reappeared within 3–7 dpa. Additionally, sensory eye cells (but not pigment cells) regenerated in juveniles at a higher proportion than in larvae (14% in juveniles, 4% in larvae). While rare, the absence of eyes immediately following amputation and the ectopic location of the observed eye cells made us confident that these cells are regenerated eye cells in amputated larvae and juveniles. One possible explanation for the variation in cilia and eye cell regeneration across individuals is variance in maternal nutritional output.

In summary, the beginning stages of regeneration (i.e., wound healing, localized cellular proliferation, and expression of stem cell markers, etc.) occur in amputated larvae but there is a lack complete replacement of lost tissue or of cell or tissue differentiation. These data indicate that *C. teleta* larvae exhibit limited regeneration abilities relative to the successful posterior regeneration observed in juvenile and adult stages. Additionally, detection of initial stages of regeneration in larvae suggests that *C. teleta* has a gradual transition in regenerative potential, rather than a stark change in potential that corresponds with the metamorphoric transition.

### 4.2 Amputation-induced expression in preexisting tissue

*MyoD* functions in muscle development and its expression in muscle progenitor cells is an indication of myogenesis (Zammit, 2017; Rawls & Olson, 1997). In response to amputation, *MyoD* expression expands across the length of the larval body in *C. teleta*, far from the wound site. This expanded expression may result in an increase in muscle cell number to compensate for a decrease in body length following amputation, although we did not observe an obvious increase in the number of muscle fibers 3 – 5 dpa. Alternatively, expanded expression of *MyoD* may be an injury-specific response that leads to cell dedifferentiation and return to a precursor state. Such a precursor cell population may contribute to regenerating tissue. It is notable that the induction of expanded *MyoD* expression occurs in the absence of a complete regeneration response. Regardless, future functional work is needed to better understand the implications of this change in *MyoD* expression in response to amputation. Changes in gene expression in response to amputation has previously been documented in *C. teleta* and other annelids (de Jong & Seaver, 2016; Kozin et al., 2017; Pfeifer et al., 2012; Takeo et al., 2008). However, most examples documented to date are changes in expression of positional identity genes, and their new expression likely reflects repatterning of the body fragment in response to amputation.

### 4.3 Metamorphosis following amputation

Amputated *C. teleta* larvae undergo successful metamorphosis. We were surprised to observe metamorphosis following anterior amputations. In other spiralians, it has been hypothesized that the brain and other anterior sensory organs function in the detection and response to metamorphic cues (Chartier et al., 2018; Conzelmann et al., 2013; Hadfield, 2010; Hadfield et al., 2000). In *C. teleta*, chemo-sensitive ciliated cells in the head ectoderm were proposed to relay metamorphosis signals directly to the cerebral ganglia (Biggers & Laufer, 1999). More recently, a study hypothesized that the inductive signal for metamorphosis is detected by sensory neurons that innervate the dorsal pharyngeal pad of the pharynx (Biggers et al., 2012). If the metamorphic cue was detected by anterior neurons, we would expect that removing the head would prevent metamorphosis. However, our results demonstrate that the brain is not required for detecting the metamorphic cue or for undergoing metamorphosis in *C. teleta*, supporting the hypothesis that detection of metamorphic cues in *C. teleta* likely occur in the pharynx. It was similarly demonstrated in the polychaete *Hydroides elegans* that apical sensory neurons are not necessary for successful metamorphosis (Nedved et al., 2021), suggesting that the brain may not be necessary for metamorphosis in some annelids.

It is yet to be determined whether the amputations performed during larval stages have lasting effects on survival and growth following metamorphosis. In addition, it remains to be seen whether juveniles induced from posteriorly amputated larvae redevelop posterior growth zones, can generate additional segments, or are capable of posterior regeneration. Future studies could examine the regeneration potential of juveniles that result from posteriorly amputated larvae. Additionally, it could be interesting to determine if juveniles induced from anterior amputated larvae are capable of posterior regeneration in the absence of a brain.

### 4.4 Larval regeneration in marine invertebrates

The regenerative potential of an animal can vary significantly across its life stages. In the literature, larvae with reduced regenerative ability relative to adults are described as having “attempted” or “limited” regeneration (Vickery et al., 2001). Using this convention, we categorize *C. teleta* larvae as having limited regeneration. Here, we show that *C. teleta* gains regeneration ability over its life cycle and exhibits different regeneration abilities between larval and juvenile stages. To our knowledge, published studies characterizing larval regeneration abilities in other annelid larvae are scarce.

Other bilaterian taxa suffer from a similar dearth in sampling larval and immature stages for their regenerative abilities. Within Spiralia, larval regeneration has been examined in one nemertean species. Similar to annelids, nemertean adult regeneration is variable (Zattara et al., 2019; Zattara & Fernández-Alvarez, 2022)*. Maculaura alaskensis* appears to have high regenerative potential as early stage larvae that progressively decreases with maturation until it stabilizes as an adult capable of posterior regeneration (Hiebert & Maslakova, 2015). Younger *M. alaskensis* larvae are five times more likely to regenerate than older larvae following amputation (Moss, 2017). In Ecdoysozoa, the other protostome clade, regeneration ability is generally restricted to appendages, rather than along the main body axis (Brenneis et al., 2023). However, recent work shows that immature instars of the sea spider *Pycnogonum littorale* can regenerate the midgut and gonads as well as appendages following bisection. Adult counterparts of *P. littorale* cannot regenerate and do not even survive the amputations, suggesting that regeneration ability is restricted to the immature instar stages (Brenneis et al., 2023).

Deuterostome larvae have been better surveyed for regeneration ability and there are numerous documented examples. The hemichordate *Ptychodera flava* has the ability to repeatedly regenerate all body parts when cut transversely or decapitated as an adult (Humphreys et al., 2022). Although *P. flava* larvae can regenerate along its main body axis, it does not regenerate as well as the adults. In addition, regeneration ability differs across the larval body. Specifically, ventral regions of *P. flava* larvae regenerate better than dorsal regions and posterior segments regenerate better than anterior segments (Luttrell et al., 2018). The larval tail of the cephalochordate *Branchistoma floridae* regenerates multiple tissues faster than adults following similar amputations (Zou et al., 2021). Unlike other clades, echinoderm larvae have been widely surveyed for regenerative potential in all classes with documented adult whole body regeneration abilities (Cary et al., 2019; Vickery et al., 2001). Asteroids and echinoid larvae, such as *Patiria miniate,* can regenerate completely, regardless of developmental stage, demonstrating a stable regenerative ability throughout life (Oulhen, 2016; Carnevali, 2006; Vickery et al., 2002). In crinoids, there appears a trend of increasing regenerative potential over developmental time. For example, crinoid larvae display limited ability to regenerate body parts, with early-stage *Antedon rosacea* larvae regenerate more poorly than late-stage larvae (Carnevali, 2006). In contrast, adults can replace lost structures even when reduced to 1/5 of the original body size (Carnevali, 2006). In summary, no clear trends of increasing or decreasing regeneration abilities across the life cycle emerge from regeneration studies either within or across taxa, suggesting changes in regeneration ability across the life cycle is a variable trait.

Although we cannot currently identify trends reflecting how regeneration abilities change between larval and adult stages from a phylogenetic perspective, we can consider the impact of life history factors. Factors such as feeding (de Jong & Seaver, 2016; Özpolat et al., 2016; Zattara & Bely, 2013), duration of larval stage, and sublethal predation (Schoeman and Simon, 2023; Stewart, 1996) may better correlate with the regenerative abilities observed in an individual species. Unlike *C. teleta*, *M. alaskensis*, *P. flava*, and *P. miniate* larvae feed and display an ability to replace complex structures following amputation. *M. alaskensis* and *P. miniate* can swim in the water column for weeks or months (Moss, 2017; Cary et al., 2019) and *P. flava* for up to 300 days (Luttrell et al., 2018). In contrast, larvae of *C. teleta* and the crinoid *A. rosacea* are non-feeding (Vickery et al., 2001) and have a short larval period before undergoing metamorphosis. *C. teleta* larvae swim in the water column for hours to a few days, and some crinoids for up to 10 days before settlement (Nakano et al., 2003). It has been previously proposed that the longer an animal is in the water column and vulnerable to predation, the greater risk there is of sublethal injury (Vickery & McClintock, 1998). Sublethal damage could act as an evolutionary pressure to select for a robust regenerative ability. To that end, there may not be a strong selective advantage for regeneration in non-feeding, short lived, brood dwelling larvae like *C. teleta*.

### 4.5 Future directions

This study demonstrates differences in regenerative ability across developmental stages of *Capitella teleta*. Future studies could investigate physiological, cellular, and molecular differences between larvae and juveniles in response to amputation. One such difference may relate to nutritional status. It has been demonstrated that the extent of regeneration in *C. teleta* juveniles and other annelids can fluctuate depending on nutrition intake (de Jong & Seaver, 2016; Zattara & Bely, 2013). Because *C. teleta* larvae are non-feeding, limited energy resources may contribute to their regeneration limitations. This nutrition hypothesis could be tested by two separate approaches. In the first, cells with high yolk content can be deleted in early-stage embryos to reduce maternally contributed nutrition. Previous work demonstrated that such deletions result in morphologically normal, albeit smaller larvae (Pernet et al., 2012). These larvae would have a reduced nutritional supply and may exhibit a further reduction in regenerative potential to what was observed in this study. The second approach would involve investigating regeneration abilities in a different annelid with a feeding larval form. A feeding larva with more nutritional resources may exhibit full regenerative abilities following amputation. Additionally, an annelid with a longer larval period would have more time to completely regenerate lost tissue following amputation during the early larval period, while also continually feeding.

The timing of cell maturation may also limit larval regeneration and explain differences between larval and juvenile regenerative abilities in *C. teleta*. An example of one such developmental constraint may be in the timing of stem cell maturation. A previous study proposed that stem cells travel from the multipotent progenitor cell (MPC) cluster to the wound site during juvenile regeneration (de Jong & Seaver, 2017). MPC cells in larvae may be immature, thereby preventing these stem cells from positively contributing to successful regeneration. Specifically, nascent stem cells may not be capable of self-renewal or migration, or of contributing to the blastema (Juliano et al., 2010). The number of cells in the MPC cluster does not increase during larval development (de Jong & Seaver, 2017), suggesting these cells may be quiescent during this time. A heterotopic transplant of a mature MPC cluster from juveniles into larvae would test this hypothesis, and an increased regenerative ability in larvae following transplantation would be expected. Such experiments in *C. teleta* regeneration might lead to a better understanding of the role that development has on regeneration.

Future studies can characterize the regeneration ability of other annelid larvae, particularly those whose adults have different regenerative potential than that of *C. teleta*. A correlation between adult and larval regenerative potential within individual species may exist. For example, there may be anterior regenerative ability in larvae in a species capable of anterior regeneration as an adult. If this correlation holds, the larvae of *Chaetopterous pergamentaceus* (Cuvier & Latreille, 1829) (formally *variopedatus*) would be predicted to regenerate both anterior and posterior structures. Adult *C. pergamentaceus* can regenerate anteriorly following amputations at segment 15 or more anterior and can regenerate posteriorly following amputation at any segment (Berrill, 1928; Cuvier & Latreille, 1829). Extensive additional sampling is needed before we can meaningfully determine whether the pattern observed in *C. teleta* is representative of other annelids. Additional sampling across annelids will also provide insight as to the extent of variation in larval regeneration abilities, and whether larval regenerative ability is as common as adult regenerative ability.

## Acknowledgements

We thank Brent Foster, Lauren Kunselman, and Katie Feerst for critical reading of the manuscript.

## Author Contributions

Conceived and designed the experiments: AAB ECS. Performed the experiments: AAB. Analyzed the data: AAB ECS. Contributed reagents/materials/analysis tools: ECS AAB. Wrote the paper: AAB ECS.

## References

Alvarado, A. S. (2000). Regeneration in the metazoans: why does it happen? BioEssays. 10.1002/(SICI)1521-1878(200006)22:6

Bely, A. E. (2006). Distribution of segment regeneration ability in the *Annelida*. Integrative and Comparative Biology, 46(4), 508–518. 10.1093/icb/icj051

Bely, A. E., & Sikes, J. M. (2010). Latent regeneration abilities persist following recent evolutionary loss in asexual annelids. Proceedings of the National Academy of Sciences of the United States of America, 107(4), 1464–1469. 10.1073/pnas.0907931107

Berrill, N.J. (1928). Regeneration in the polychaet *Chaetopterus variopedatus*. Journal of Marine Biological Association of the United Kingdom, 15(1), 151–158. 10.1017/S0025315400055594

Biggers, W. J., & Laufer, H. (1999). Settlement and metamorphosis of Capitella larvae induced by juvenile hormone-active compounds is mediated by protein kinase C and ion channels. 196(2), 187–198. 10.2307/1542564

Biggers, W. J., Pires, A., Pechenik, J. A., Johns, E., Patel, P., Polson, T., & Polson, J. (2012). Inhibitors of nitric oxide synthase induce larval settlement and metamorphosis of the polychaete annelid *Capitella teleta*. Invertebrate Reproduction and Development, 56(1), 1–13. 10.1080/07924259.2011.588006

Blake, J.A., Grassle, J.P., & Eckelbarger, K.J. (2009). *Capitella teleta*, a new species designation for the opportunistic and experimental *Capitella* sp. I, with a review of the literature for confirmed records. Zoosymposia, 2(1), 25– 53. DOI:10.11646/zoosymposia.2.1.6

Boyle, M. J., & Seaver, E. C. (2008). Developmental expression of *foxA* and *gata* genes during gut formation in the polychaete annelid, Capitella sp. I. Evolution & Development, 10(1), 89–105. 10.1111/J.1525-142X.2007.00216.X

Brenneis, G., Frankowski, K., MaaB, L., & Scholtz, G. (2023) The sea spider *Pycnogonum littorale* overturns the paradigm of the absence of axial regeneration in molting animals. PNAS, 120(5). 10.1073/pnas.221727120

Brockes, J. P. (1997). Amphibian limb regeneration: rebuilding a complex structure. Science, 276(5309), 81–87. 10.1126/SCIENCE.276.5309.81

Burns, R. T., Pechenik, J. A., Biggers, W. J., Scavo, G., & Lehman, C. (2014). The B vitamins nicotinamide (B3) and riboflavin (B2) stimulate metamorphosis in larvae of the deposit-feeding polychaete capitella teleta: implications for a sensory ligand-gated ion channel. PLoS ONE, 9(11). 10.1371/journal.pone.0109535

Byrnes, E. F. (1904). Regeneration of the anterior limbs in the tadpoles of frogs. Archiv Für Entwicklungsmechanik Der Organismen, 18(2), 171–177. 10.1007/BF02163652

Carnevali, C. (2006). Regeneration in echinoderms: repair, regrowth, cloning. Invertebrate Survival Journal. 3(1), 64–76.

Carnevali, M. D. C., Lucca, E., & Bonasoro, F. (1993). Mechanisms of arm regeneration in the feather star *Antedon mediterranea*: healing of wound and early stages of development. Journal of Experimental Zoology, 267(3), 299–317. 10.1002/JEZ.1402670308

Cary, G. A., Wolff, A., Zueva, O., Pattinato, J., & Hinman, V. F. (2019). Analysis of sea star larval regeneration reveals conserved processes of whole-body regeneration across the metazoa. BMC Biology, 17(1), 16. 10.1186/s12915-019-0633-9

Chartier, T. F., Deschamps, J., Dürichen, W., Jékely, G., & Arendt, D. (2018). Whole-head recording of chemosensory activity in the marine annelid *Platynereis dumerilii*. Open Biology, 8(10). 10.1098/RSOB.180139

Cohen, R. A., & Pechenik, J. A. (1999). Relationship between sediment organic content, metamorphosis, and postlarval performance in the deposit-feeding polychaete *Capitella* sp. I Journal of Experimental Marine Biology and Ecology, 240, 1–18. 10.1016/S0022-0981(99)00047-7

Conzelmann, M., Williams, E. A., Tunaru, S., Randel, N., Shahidi, R., Asadulina, A., Berger, J., Offermanns, S., & Jékely, G. (2013). Conserved MIP receptor-ligand pair regulates *Platynereis* larval settlement. Proceedings of the National Academy of Sciences of the United States of America, 110(20), 8224– 8229. 10.1073/PNAS.1220285110/-/DCSUPPLEMENTAL

Cuvier, G.*, &* Latreille, P. A. (1829). Le règne animal distribué d’après son organisation, pour servir de base à l’histoire naturelle des animaux et d’introduction à l’anatomie comparée. Chez Déterville. 10.5962/bhl.title.49223

Dannenberg, L. C., & Seaver, E. C. (2018). Regeneration of the germline in the annelid *Capitella teleta*. Developmental Biology, 440(2), 74–87. 10.1016/j.ydbio.2018.05.004

de Jong, D. M., & Seaver, E. C. (2016). A stable thoracic hox code and epimorphosis characterize posterior regeneration in *Capitella teleta*. PLOS ONE, 11(2), e0149724. 10.1371/journal.pone.0149724

de Jong, D. M., & Seaver, E. C. (2017). Investigation into the cellular origins of posterior regeneration in the annelid *Capitella teleta*. *Regeneration (Oxford*, England), 5(1), 61–77. 10.1002/reg2.94

del Olmo, I., Verdes, A., & Alvarez-Campos, P. (2022) Distinct patterns of gene expression during regeneration and asexual reproduction in the annelid *Pristina leidyi*. Journal of experimental zoology. doi: 10.1002/jez.b.23143

Dill, K. K., & Seaver, E. C. (2008). *Vasa* and *nanos* are coexpressed in somatic and germ line tissue from early embryonic cleavage stages through adulthood in the polychaete *Capitella* sp. I. Development Genes and Evolution, 218(9), 453–463. 10.1007/s00427-008-0236-x

Giani, V. C., Yamaguchi, E., Boyle, M. J., & Seaver, E. C. (2011). Somatic and germline expression of *piwi* during development and regeneration in the marine polychaete annelid *Capitella teleta*. EvoDevo, 2(1). 10.1186/2041-9139-2-10

c. Good Calculators. [online software]. https://goodcalculators.com/one-way-anova-calculator/

Grassle, J., & Grassle, J. F. (1976). Sibling species in the marine pollution indicator *Capitella* (polychaeta). Science 192(4239),567– 569. DOI: 10.1126/science.1257794

Hadfield, M. G. (2010). Biofilms and marine invertebrate larvae: What bacteria produce that larvae use to choose settlement sites. Annual review of marine science, 3, 453–470. 10.1146/ANNUREV-MARINE-120709-142753

Hadfield, M. G., Meleshkevitch, E. A., & Boudko, D. Y. (2000). The apical sensory organ of a gastropod veliger is a receptor for settlement cues. Biological Bulletin, 198(1):67–76. doi: 10.2307/1542804

Harrison, R. G. (1898). The growth and regeneration of the tail of the frog larva – Studied with the aid of Born’s method of grafting. Archiv Für Entwickelungsmechanik Der Organismen, 7(2–3), 430–485. 10.1007/BF02161494

Hiebert, T. C., & Maslakova, S. (2015). Integrative taxonomy of the Micrura alaskensis Coe, 1901 species complex (Nemertea: Heteronemertea), with descriptions of a new genus maculaura gen. nov. and four new species from the NE pacific. Zoological Science, 32(6), 615–637. 10.2108/ZS150011

Humphreys, T., Weiser, K., Arimoto, A., Sasaki, A., Uenishi, G., Fujimoto, B., Kawashima, T., Taparra, K., Molnar, J., Satoh, N., Marikawa, Y., & Tagawa, K. (2022) Ancestral stem cell reprogramming genes active in hemichordate regeneration. Frontiers in Ecology and Evolution. doi:10.3389/fevo.2022.769433

Jeffery, W. R. (2015). Regeneration, stem cells, and aging in the tunicate *Ciona*: insights from the oral siphon. International Review of Cell and Molecular Biology, 319, 255–282. 10.1016/bs.ircmb.2015.06.005

Juliano, C. E., Swartz, S. Z., & Wessel, G. M. (2010). A conserved germline multipotency program. Development, 137(24), 4113–4126. 10.1242/DEV.047969

Kostyunchenko, R.P. (2022) *Nanos* is expressed in somatic and germline tissue during larval and post-larval development of the annelid *Alitta virens*. Genes.13, 270. http://doi.org/10.3390/genes13020270

Lowry, R. (2023) vassarstats [online software]. http://vassarstats.net/anova1u.html

Luttrell, S. M., Su, Y. H., & Swalla, B. J. (2018). Getting a head with *Ptychodera flava* larval regeneration. Biological Bulletin, 234(3), 152–164. 10.1086/698510

Meyer, N. P., Boyle, M. J., Martindale, M. Q., & Seaver, E. C. (2010). A comprehensive fate map by intracellular injection of identified blastomeres in the marine polychaete *Capitella teleta*. EvoDevo, 1(1). 10.1186/2041-9139-1-8

Meyer, N. P., Carrillo-Baltodano, A., Moore, R. E., & Seaver, E. C. (2015). Nervous system development in lecithotrophic larval and juvenile stages of the annelid *Capitella teleta*. Frontiers in Zoology, 12(1), 1–27. 10.1186/s12983-015-0108-y

Meyer, N. P., & Seaver, E. C. (2009). Neurogenesis in an annelid: Characterization of brain neural precursors in the polychaete *Capitella* sp. I. Developmental Biology, 335(1), 237–252. 10.1016/J.YDBIO.2009.06.017

Nakano, H., Hibino, T., Oji, T., Hara, Y., & Amemiya, S. (2003) Larval stages of a living sea lily (stalked crinoid echinoderm). Nature 421, 158–160. 10.1038/nature01236

Nedved, B. T., Freckelton, M. L., & Hadfield, M. G. (2021). Laser ablation of the apical sensory organ of *Hydroides elegans* (Polychaeta) does not inhibit detection of metamorphic cues. The Journal of Experimental Biology, 224(20). 10.1242/JEB.242300

Oulhen, N., Heyland, A., Carrier, T. J., Zazueta-Novoa, V., Fresques, T., Laird, J., Onorato, T. M., Janies, D., & Wessel, G. (2016). Regeneration in bipinnaria larvae of the bat star *Patiria miniata* induces rapid and broad new gene expression. Mechanisms of Development, 142, 10–21. 10.1016/j.mod.2016.08.003

Özpolat, B. D., & Bely, A. E. (2016). Developmental and molecular biology of annelid regeneration: a comparative review of recent studies. Current Opinion in Genetics & Development, 40, 144–153. 10.1016/j.gde.2016.07.010

Özpolat, B.D., Sloane, E.S., Zattara, E.E., and Bely, A.E. (2016) Plasticity and regeneration of gonads in the annelid *Pristina leidyi*. EvoDevo 7 (22). 10.1186/s13227-016-0059-1

Pernet, B., Amiel, A., & Seaver, E. C. (2012). Effects of maternal investment on larvae and juveniles of the annelid *Capitella teleta* determined by experimental reduction of embryo energy content. Invertebrate Biology, 131(2), 82–95. 10.1111/j.1744-7410.2012.00263.x

Phipps, L. S., Marshall, L., Dorey, K., & Amaya, E. (2020). Model systems for regeneration: *Xenopus*. Development (Cambridge), 147(6). 10.1242/DEV.180844/223048

Planques, A., Malem, J., Parapar, J., Vervoot, M., & Gazave, E. (2019) Morphological, cellular and molecular characterization of posterior regeneration in the marine annelid *Playnereis dummerilli*. Developmental Biology 445(189-210). http://doi.org/10.10106/j.ydbio.2018.11.004

Rawls, A., & Olson, E. N. (1997). *MyoD* meets its maker. Cell, 89(1), 5–8. 10.1016/S0092-8674(00)80175-0

Schindelin, J., Arganda-Carreras, I., Frise, E., Kaynig, V., Longair, M., Pietzsch, T., Preibisch, S., Rueden, C., Saalfeld, S., Schmid, B., Tinevez, J. Y., White, D. J., Hartenstein, V., Eliceiri, K., Tomancak, P., & Cardona, A. (2012). Fiji: an open-source platform for biological-image analysis. Nature Methods 2012 9:7, 9(7), 676–682. 10.1038/nmeth.2019

Schoeman, S., & Simon, C.A. (2023). Live to die another day: regeneration in Diopatra aciculata Knox and Cameron, 1971 (Annelida: Onuphidae) collected as bait in Knysna Estuary, South Africa. Biology. 12(3), 483. 10.3390/biology12030483

Seaver, E. C. (2022). Sifting through the mud: a tale of building the annelid *Capitella teleta* for EvoDevo studies. Current Topics in Developmental Biology, 147, 401–432. 10.1016/BS.CTDB.2021.12.018

Seaver, E. C., Thamm, K., & Hill, S. D. (2005). Growth patterns during segmentation in the two polychaete annelids, Capitella sp. I and Hydroides elegans: comparisons at distinct life history stages. Evolution Development, 7(4), 312–326. 10.1111/j.1525-142X.2005.05037.x

Singer, M. (1954). Induction of regeneration of the forelimb of the postmetamorphic frog by augmentation of the nerve supply. Journal of Experimental Zoology, 126(3), 419–471. 10.1002/jez.1401260304

Stewart, B. (1998). Sub-lethal predation and rate of regeneration in the euryalinid snake star *Astrobrachion constrictum* (Echinodermata, Ophiuroidea) in a New Zealand fiord. Journal of Experimental Marine Biology and Ecology. 199(2), 269–283. 10.1016/0022-0981(95)00177-8

Sur, A., Magie, C. R., Seaver, E. C., & Meyer, N. P. (2017). Spatiotemporal regulation of nervous system development in the annelid Capitella teleta. 8, 13. 10.1186/s13227-017-0076-8

Vickery, M.S., McClinktock, J.B. (1998). Regeneration in metazoan larvae. Nature. 394, 140. 10.1038/28086

Vickery, M. C. L., Vickery, M. S., Amsler, C. D., & McClintock, J. B. (2001). Regeneration in echinoderm larvae. Microscopy Research and Technique, 55(6), 464–473. 10.1002/jemt.1191

Vickery, Minako S., Vickery, M. C. L., & McClintock, J. B. (2002). Morphogenesis and organogenesis in the regenerating planktotrophic larvae of asteroids and echinoids. Biological Bulletin, 203(2), 121–133. 10.2307/1543381

Wilhelm, E., Bückmann, D., & Tomaschko, K.H. (1997). Life cycle and population dynamics of *Pycnogonum litorale* (Pycnogonida) in a natural habitat. Marine Biology 129, 601–606. 10.1007/s002270050202

Yamaguchi, E., Dannenberg, L. C., Amiel, A. R., & Seaver, E. C. (2016). Regulative capacity for eye formation by first quartet micromeres of the polychaete *Capitella teleta*. Developmental Biology, 410(1), 119–130. 10.1016/j.ydbio.2015.12.009

Yamaguchi, E., & Seaver, E. C. (2013). The importance of larval eyes in the polychaete *Capitella teleta*: effects of larval eye deletion on formation of the adult eye. Invertebrate Biology, 132(4), 352–367. 10.1111/ivb.12034

Zammit, P. S. (2017). Function of the myogenic regulatory factors Myf5, MyoD, Myogenin and MRF4 in skeletal muscle, satellite cells and regenerative myogenesis. Seminars in Cell & Developmental Biology, 72, 19–32. 10.1016/J.SEMCDB.2017.11.011

Zattara, E. E., & Bely, A. E. (2013). Investment choices in post-embryonic development: quantifying interactions among growth, regeneration, and asexual reproduction in the annelid *Pristina leidyi*. Journal of Experimental Zoology Part B: Molecular and Developmental Evolution, 320(8), 471–488. 10.1002/jez.b.22523

Zattara, E. E., & Bely, A. E. (2016). Phylogenetic distribution of regeneration and asexual reproduction in *Annelida*: regeneration is ancestral and fission evolves in regenerative clades. Invertebrate Biology, 135(4), 400–414. 10.1111/IVB.12151

Zattara, E.E. & Fernández-Alvarez, F.A. (2022) Collecting and culturing *Lineus sanguineus* to study *Nemertea* WBR. In: Blanchoud S, Galliot B, editors. Whole-Body Regeneration: Methods and Protocols [Internet]. New York (NY) Chapter 12. doi: 10.1007/978-1-0716-2172-1_12

Zattara E.E., Fernández-Álvarez F. A., Hiebert T. C., Bely A. E. & Norenburg J. L. (2019) A phylum-wide survey reveals multiple independent gains of head regeneration in Nemertea. The Royal Society. 10.1098/rspb.2018.2524

Zou, J., Wu, X., Shi, C., Zhong, Y., Yan, Q., Su, L., and Li, G. (2021) A potential method for rapid screening of amphioxus founder harboring germline mutation and transgene. Front. Cell Dev. Biol. 9 (702290) doi: 10.3389/fcell.2021.702290

